# SCDC: Bulk Gene Expression Deconvolution by Multiple Single-Cell RNA Sequencing References

**DOI:** 10.1101/743591

**Authors:** Meichen Dong, Aatish Thennavan, Eugene Urrutia, Yun Li, Charles M. Perou, Fei Zou, Yuchao Jiang

## Abstract

Recent advances in single-cell RNA sequencing (scRNA-seq) enable characterization of transcriptomic profiles with single-cell resolution and circumvent averaging artifacts associated with traditional bulk RNA sequencing (RNA-seq) data. Here, we propose SCDC, a deconvolution method for bulk RNA-seq that leverages cell-type specific gene expression profiles from multiple scRNA-seq reference datasets. SCDC adopts an ENSEMBLE method to integrate deconvolution results from different scRNA-seq datasets that are produced in different laboratories and at different times, implicitly addressing the problem of batch-effect confounding. SCDC is benchmarked against existing methods using both *in silico* generated pseudo-bulk samples and experimentally mixed cell lines, whose known cell-type compositions serve as ground truths. We show that SCDC outperforms existing methods with improved accuracy of cell-type decomposition under both settings. To illustrate how the ENSEMBLE framework performs in complex tissues under different scenarios, we further apply our method to a human pancreatic islet dataset and a mouse mammary gland dataset. SCDC returns results that are more consistent with experimental designs and that reproduce more significant associations between cell-type proportions and measured phenotypes.

## Introduction

Bulk RNA sequencing (RNA-seq) has been the method of choice for profiling transcriptomic variations under different conditions such as disease states (Robinson et al., 2010, Love et al., 2014, Ritchie et al., 2015). However, in complex tissues with multiple heterogeneous cell types, bulk RNA-seq measures the average gene expression levels by summing over the population of cells in the tissue, and variability in cell-type compositions confounds with analysis such as detecting differential gene expression (Avila Cobos et al., 2018). While multiple statistical and computational methods have been developed for cell-type decomposition of bulk RNA-seq data (Shen-Orr et al., 2010, Gong and Szustakowski 2013, Newman et al., 2015), most of these have limitations. Many require a priori knowledge, either of gene expression profiles of purified cell types (Gong and Szustakowski 2013, Newman et al., 2015) or of cell-type compositions (Shen-Orr et al., 2010). Methods that do not take these information as input instead require a list of pre-selected marker genes (Zhong et al., 2013, Becht et al., 2016). Finally, completely unsupervised approaches based on non-negative matrix factorization suffer from low deconvolution accuracy and have identifiability and multicollinearity issues (Wang et al., 2014).

Recent advances in single-cell RNA sequencing (scRNA-seq) circumvent averaging artifacts associated with the traditional bulk RNA-seq data by enabling characterization of transcriptomic profiles at the single-cell level (Saliba et al., 2014). While scRNA-seq data has greatly increased resolution in the characterization of transcriptomic heterogeneity, its relatively high cost and technical challenges pose difficulties in generating scRNA-seq data across a large population of samples (Stegle et al., 2015, Ziegenhain et al., 2017). Association testing performed on single-cell data from a small number of subjects has only limited statistical power. Large collaborations, on the other hand, have successfully sequenced an enormous number of bulk samples (Edgar et al., 2002, National Cancer Institute, 2019), making cell-type decomposition on bulk RNA-seq data aided by scRNA-seq an appealing analysis scheme.

Several methods exploiting single-cell expression reference datasets have been developed for bulk gene expression deconvolution (Baron et al., 2016, Wang et al., 2019, Newman et al., 2019, Jew et al., 2019). Specifically, Bseq-SC (Baron et al., 2016) uses scRNA-seq data to build a cell-type specific gene expression signature matrix for a set of pre-selected marker genes, then applies a support vector regression-based deconvolution framework adapted from CIBERSORT (Newman et al., 2015). Similarly, Bisque (Jew et al., 2019) and CIBERSORTx (Newman et al., 2019) take as input a list of pre-selected marker genes and explicitly account for the technical variation in the generation of the single-cell signature matrix and the observed bulk expression. MuSiC (Wang et al., 2019) proposes a weighted non-negative least squares (W-NNLS) regression framework to utilize all genes that are shared between the bulk and the single-cell data. Genes are weighted by cross-subject and cross-cell variations and empirical evidence suggests that this leads to higher deconvolution accuracy.

Despite this progress, to the best of our knowledge, all existing methods reconstruct the gene expression signature matrix using only one single-cell reference. These methods therefore cannot use additional scRNA-seq data of the same tissue from the same model organism that may be available from other studies and laboratories (Table S1 and Figure S1). These methods also cannot take advantage of the extensive transcriptomic reference maps at the cellular level that have been generated by multiple large consortia, including the Human Cell Atlas (*Human Cell Atlas*, 2019) and the Mouse Cell Atlas (Mouse *Cell Atlas*, 2019). Borrowing information from existing data could potentially boost the performance of and increase the robustness of deconvolution. This has been demonstrated by Vallania et al. (2018), who showed that leveraging heterogeneity across multiple reference datasets could increase deconvolution accuracy and reduce biological and technical biases for microarray data. For scRNA-seq data, however, significant batch effect prevails across data collected from different sources and as we demonstrate later, the naive pooling of multiple scRNA-seq datasets to build a “mega” reference profile performs poorly. One potential solution is to correct for the batch effect in the data. However, existing batch correction methods for scRNA-seq data either adopt a dimension reduction technique for visualization and clustering (Butler et al., 2018) or change the scale of the original gene expression measurements (Haghverdi et al., 2018), both of which make subsequent deconvolution difficult-perhaps even infeasible.

Here, we introduce a new framework, SCDC, to leverage multiple ***S***ingle-***C***ell RNA-seq reference sets for bulk gene expression ***D***e***C***onvolution. Specifically, when multiple scRNA-seq reference sets are available, SCDC adopts an ENSEMBLE method to integrate deconvolution results across datasets; it implicitly addresses the problem of batch-effect confounding by giving higher weights to the scRNA-seq data that are more closely related to the bulk RNA-seq data. We benchmark our method against existing methods using pseudo-bulk samples generated *in silico*, whose true underlying cell type identities are known. We also evaluate the performance of SCDC on an RNA-seq dataset of paired single cells and bulk samples, the latter of which have experimentally controlled cell-type proportions as ground truths. SCDC is shown to out-perform existing methods by integrating multiple scRNA-seq datasets; even with only one single-cell dataset, SCDC yields enhanced deconvolution accuracy. To further demonstrate the ENSEMBLE method, SCDC is applied to two real datasets, human pancreatic islets and mouse mammary glands, using multiple scRNA-seq inputs. We show that, compared to existing methods, SCDC returns results that are more consistent with experimental designs and that reproduce more significant associations between cell-type proportions and measured phenotypes. SCDC is available as an open-source R package at https://github.com/meichendong/SCDC.

## Results

### Overview of SCDC and Deconvolution via ENSEMBLE

Figure 1 gives an overview of SCDC. The same set of bulk RNA-seq samples can be deconvoluted using different single-cell reference datasets. Empirically, we show that this may return distinct cell-type proportion estimations, due to both intrinsic biological variation and technical noise (Table S1) (Jiang et al., 2017). It is further shown that naively pooling all available single cells from different sources suffers from the prevalent batch effects and the biological heterogeneity that are present in the data (Table S1). To resolve this discrepancy while making full use of all available scRNA-seq reference datasets, SCDC adopts an ENSEMBLE method to combine the deconvolution results from individual datasets. The weights for each dataset are selected via optimization, with higher weights assigned to single-cell reference datasets that better recapitulate the true underlying gene expression profiles of the bulk samples.

**Figure 1.**
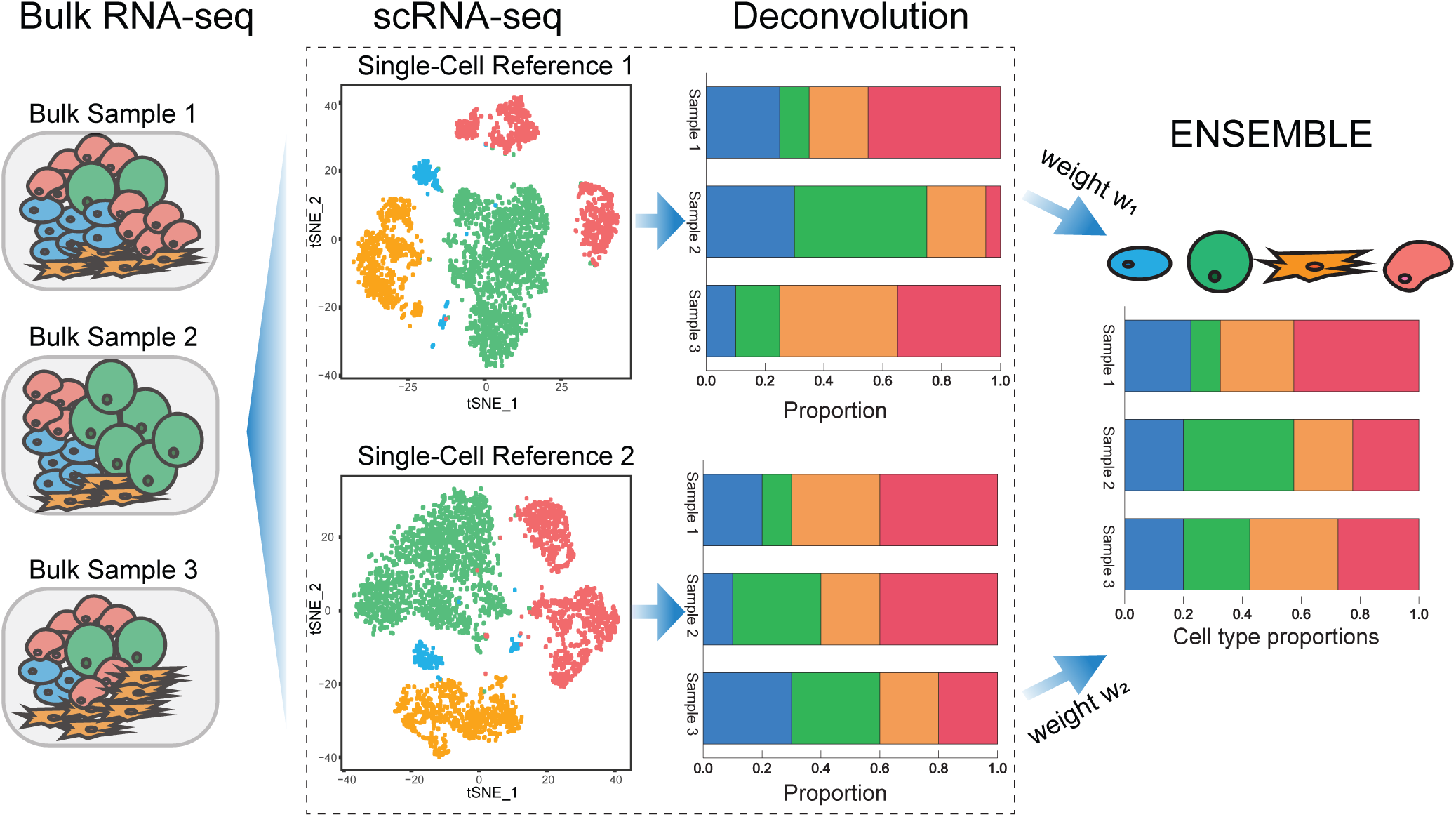
Overview of deconvolution via ENSEMBLE by SCDC. When multiple single-cell reference datasets are available, batch effect confounding is avoided by performing deconvolution on each scRNA-seq reference set separately. SCDC then integrates the deconvolution results with dataset-specific optimized weights, which are used to derive the final cell-type proportions.

In the following, we begin by giving a review of the existing regression-based deconvolution framework (Baron et al., 2016, Wang et al., 2019, Newman et al., 2019, Jew et al., 2019). We then describe the model for SCDC, leaving algorithmic details to the Methods section and Supplemental Information. Consider an observed bulk gene expression matrix **Y** ∈ ℝ ^*N×M*^ for *N* genes across *M* samples, each containing *K* different cell types. The goal of deconvolution is to find two non-negative matrices **B** ∈ ℝ^*N×K*^ and **P** ∈ ℝ^*K×M*^ such that

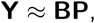

where each column of **P** represents the mixing proportions of the *K* cell types of one sample, and each column of the “basis” matrix **B** represents the average gene expression levels in each type of cells. As described earlier, different methods have been developed to integrate both bulk-tissue and single-cell gene expression measurements for deconvolution (Baron et al., 2016, Wang et al., 2019, Newman et al., 2019, Jew et al., 2019). These methods obtain:

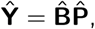

where each matrix is estimated as the final output.

In the presence of multiple scRNA-seq datasets, one can adopt the aforementioned deconvolution strategies and apply them to each single-cell dataset *r* ∈ {1, …. *R*} separately to obtain the predicted gene expression level **Ŷ**_*r*_, the estimated basis matrix 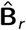, and the estimated cell-type proportion matrix 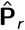. Here, **{B**_1_, **B**_2_, …, **B**_*R*_} are assumed to come from the same distribution, but with variation that arises from both the technical batch effect and from biological heterogeneity. Empirical evidence suggests that, depending on the scRNA-seq data adopted,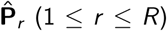 from the *R* reference datasets can differ drastically and that naively pooling all the single-cell data to estimate 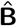 could lead to the worst performance overall (Table S1). To make full use of all available single-cell data and to give higher weights to the reference that more closely recapitulates the true underlying cell-specific gene expression profiles, SCDC adopts an ENSEMBLE method to integrate all deconvolution results with different weights *ŵ*_*r*_ (1 ≤ *r ≤ R*), which are optimized via:

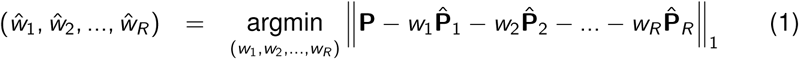

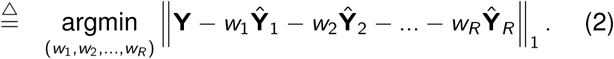

In an ideal situation, one where we know the actual cell-type proportions **P**, we would minimize the difference between the linearly weighted cell-type proportion estimates 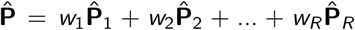 and the actual proportions **P.** However, in real dataset analysis, we do not have the luxury of a priori knowledge on the underlying **P.** Therefore, SCDC adopts a “surrogate” metric on the observed **Y** to substitute on the unknown **P.** That is, we minimize the difference between the predicted gene expressions **Ŷ** =*w*_1_**Ŷ**1 +*w*_2_ **Ŷ**_2_ + … + *W*_*R*_**Ŷ**_*R*_ and the observed gene expressions **Y.** Empirically, we show that the estimation errors on **P** are positively correlated with those on **Y** (Figure S1). That is, a reference set that leads to higher deconvolution accuracy also has lower residuals of **Y** from the regression. We also show that the *L*1 norm of the difference in equation (1) can be replaced by other dissimilarity measurements such as *L*2 norm of the difference or correlation (Figure S1). For optimization of weights (*w*_1_, …, *w*_*R*_), SCDC, by default, adopts a numerical method based on grid search.

### Performance on Simulated Data

To assess the performance of SCDC, we carried out extensive simulation studies, which also illustrate the ENSEMBLE method by SCDC in more details. In these simulations, pseudo-bulk samples were generated *in silico* by aggregating well-characterized single cells from existing scRNA-seq studies. The known cell-type proportions of these samples were used as ground truths, and the deconvolution accuracy was assessed by Pearson correlation, mean absolute deviation (mAD), and root mean square deviation (RMSD) between the actual and the deconvoluted cell-type proportions. Figure 2A gives an outline of the simulation setup. We started with a scenario where bulk RNA-seq data was paired with scRNA-seq data generated from the same study on the same subjects (Figure 2B). We then moved onto a more difficult case where the bulk RNA-seq data was generated from a different source than the scRNA-seq data (Figure 2C).

**Figure 2.**
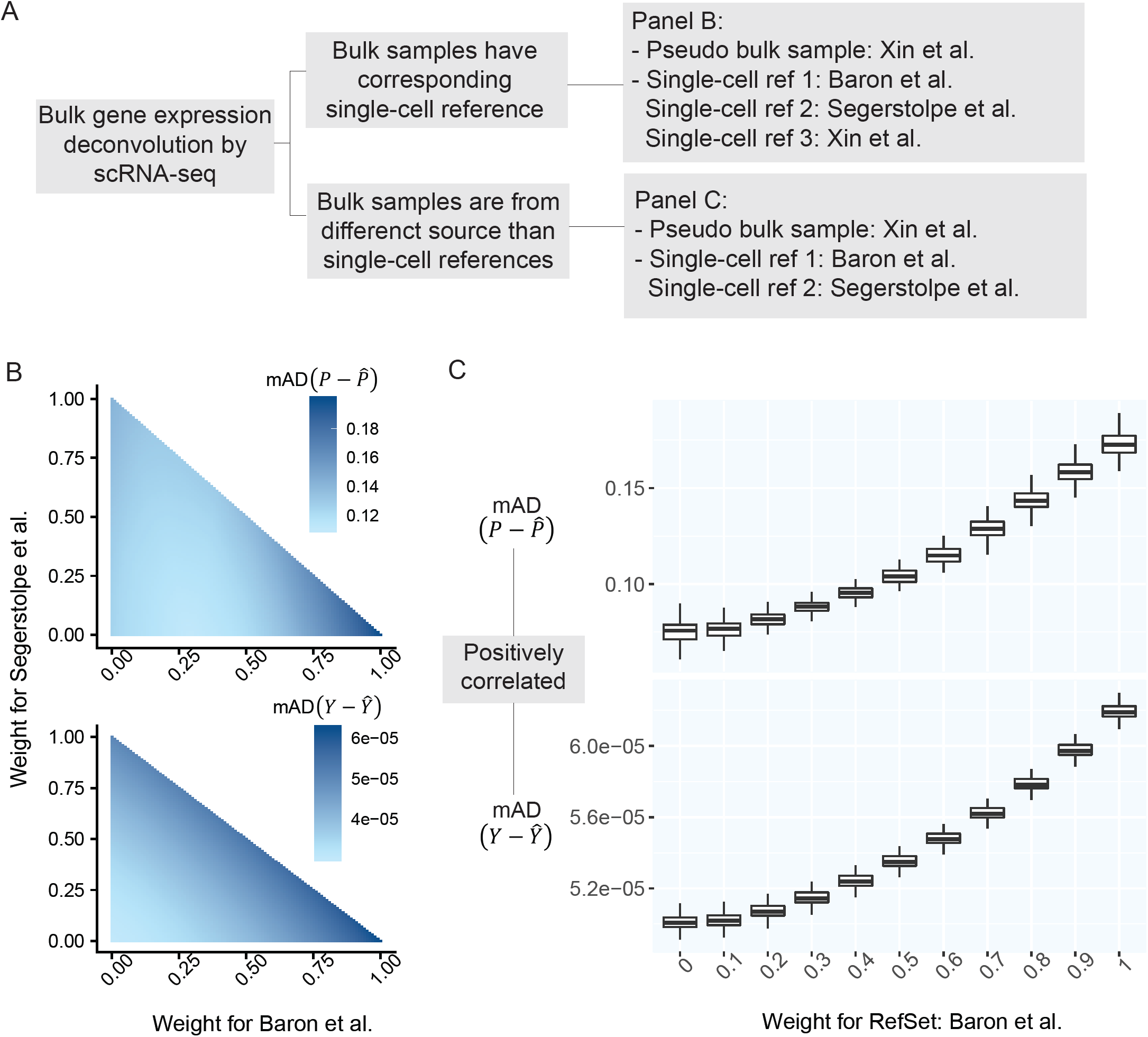
Prediction errors of Y serve as a surrogate for the estimation errors of P. **A:** Outline of simulation setup, where single cells of human pancreatic islets are aggregated to generate pseudo-bulk samples, whose cell-type proportions are known. We examine the results of deconvolution via ENSEMBLE, both with and without paired single-cell reference dataset. 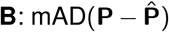 and mAD(**Y − Ŷ**) with three varying dataset-specific weights for deconvolution of bulk samples with paired scRNA-seq. The two metrics agreed on the assignment of the optimal weights, which were around (*ŵ*_1_, *ŵ*_2_, *ŵ*_3_) = (0, 0, 1).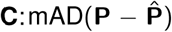 and mAD(**Y − Ŷ**) with two varying dataset-specific weights for deconvolution of bulk samples without paired scRNA-seq. The two metrics are highly correlated with varying weights for reference dataset from Baron et al. (2016).

In Figure 2B, pseudo-bulk samples were constructed by aggregating well characterized single cells of four cell types (human pancreatic alpha, beta, delta, and gamma cells) from Xin et al. (2016). 100 simulations were run. Within each run, 100 pseudo-bulk samples were generated by sampling single cells without replacement from a randomly selected subject. For deconvolution, we further adopted three scRNA-seq datasets of human pancreatic islets: Baron et al. (2016), Åsa Segerstolpe et al. (2016), Xin et al. (2016), the last of which is from the same source as the pseudo-bulk samples. In Figure 2B, we demonstrate how different weights for the three scRNA-seq reference sets (only two weights are shown since the three sum up to one) lead to different deconvolution results/accuracies, as measured by the mAD of 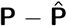 (top panel) and the mAD of **Y** -**Ŷ** (bottom panel), respectively. We show that the two metrics, given varying weights for the three single-cell reference datasets, are highly correlated, indicating that the measurement error of **Y** serves as a good proxy to that of **P.** This signifies the feasibility of the ENSEMBLE framework by SCDC when the true underlying **P** remains unknown. Indeed, our findings further reveal that SCDC was able to derive a set of optimal weights with the highest one being close to one, which corresponds to the single-cell data from the same source as the bulk samples. The same pattern is observed when we switch the source of the pseudo-bulk samples (Figure S2).

Figure 2C shows results from another set of simulations. These simulations are similar to the previously described set, but there was no scRNA-seq reference set from the same source as the pseudo-bulk samples. For pseudo-bulk samples generated from Baron et al. (2016) and Xin et al. (2016), the scRNA-seq dataset from Åsa Segerstolpe et al. (2016) is weighted most heavily by SCDC (Figure 2C, Figure S2C), potentially due to the high sequencing depth and full-transcript coverage by the Smart-seq2 protocol (Picelli et al., 2014) that was adopted. Interestingly, for the pseudo-bulk samples generated from Åsa Segerstolpe et al. (2016), SCDC recommends using weighted results from the two reference datasets (Figure S2F), highlighting the utility of the ENSEMBLE method.

### Performance on Real Dataset #1: Mixtures of Three Cell Lines with Known Proportions

While we have successfully demonstrated that SCDC allows accurate deconvolution of pseudo-bulk samples, the *in silico* reconstruction procedure is over simplified and does not mimic how real bulk RNA-seq samples are collected and sequenced. Therefore, we carried out a set of well controlled experiments, where cell lines were mixed at a fixed ratio, followed by both bulk and single-cell RNA-seq. These known cell-type proportions served as ground truths to benchmark SCDC against existing methods without bias. Specifically, human breast cancer cell lines MDA-MB-468, MCF-7, and human fibroblast cells were independently cultured and then mixed at a fixed ratio of 6:3:1. This was followed by traditional bulk RNA-seq as well as scRNA-seq by 10X Genomics. More experimental details are available in the Methods section. Single-cell clustering was performed using the Seurat pipeline (Butler et al., 2018) with t-SNE visualization shown in Figure 3A (see details in Supplemental Information). The cell-type ratio by scRNA-seq is 0.661:0.225:0.114, close to but slightly different from the experimental setup due to either the inaccuracy of counting cells when making the mixture or the sampling bias of scRNA-seq.

**Figure 3.**
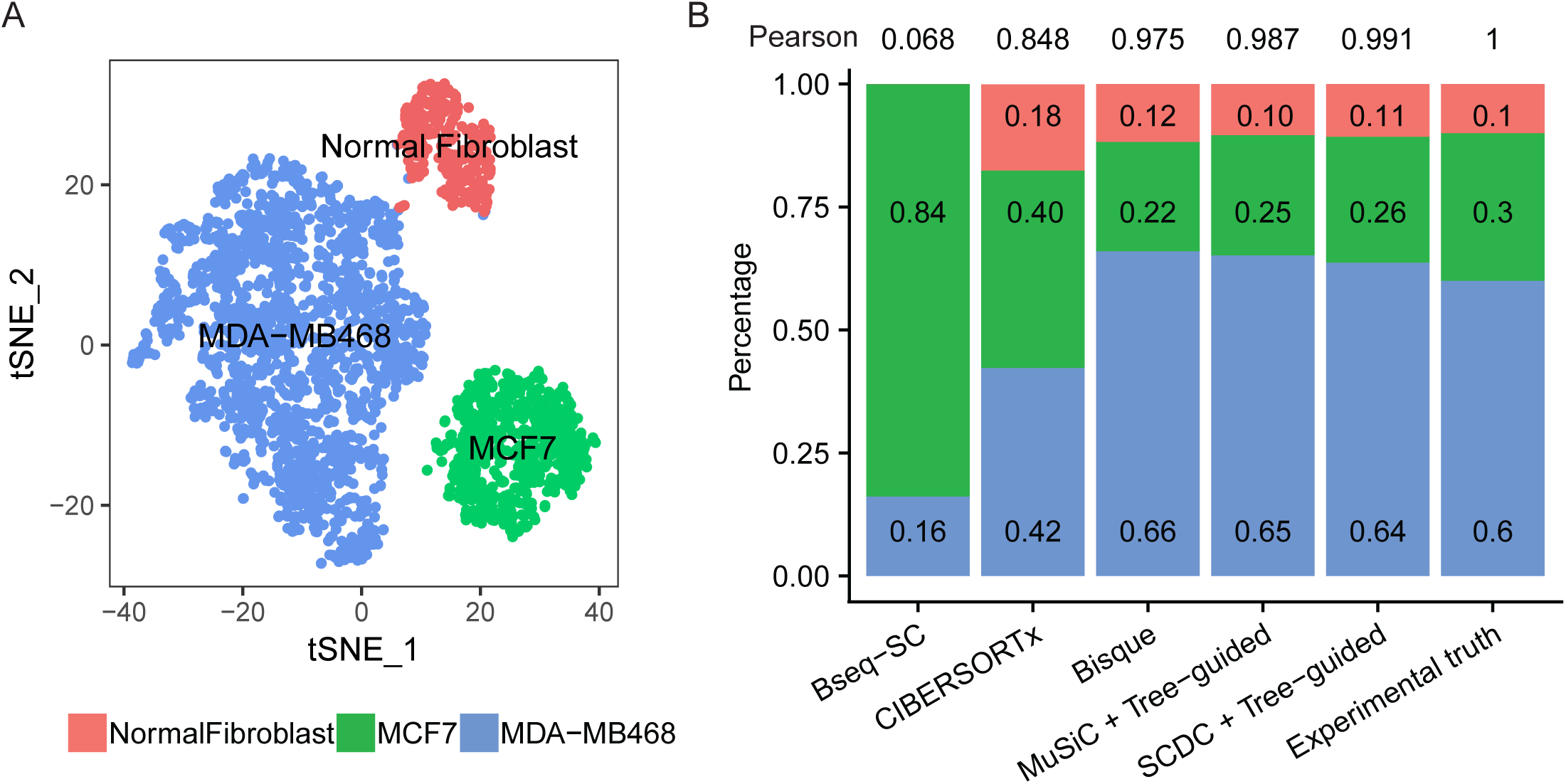
Performance assessment on bulk and single-cell RNA-seq of cell line mixtures with experimentally controlled proportions. **A:** Visualization by t-SNE after single-cell clustering. The cells are clustered into three groups, MDA-MB-468, MCF-7, and normal fibroblast cells, in a ratio close to 6:3:1. **B:** Benchmark of deconvolution results for the bulk RNA-seq sample, produced by different methods. Among all benchmarked methods, the proportions estimated by SCDC using the tree-guided approach has the highest Pearson correlation (0.99) with the ground truth.

To deconvolute the bulk RNA-seq sample, we adopted the scRNA-seq dataset that was generated from the same mixture, which was the only available reference set. As such, this reduced to a one-subject and one-reference deconvolution problem (see Supplemental Information for details), and the ENSEMBLE step was therefore not needed. In this case, we carried out direct comparisons of SCDC without ENSEMBLE against existing methods. Given one single-cell reference dataset, SCDC largely follows the W-NNLS framework proposed by MuSiC but also differs in several ways. First, SCDC starts by scaling the raw single-cell read-count matrix by a gene-and subject-specific maximal variance weight so that residuals from genes with larger weights have smaller impact on cell-type composition estimation. Second, SCDC does not take cell-type memberships as granted; instead it removes potentially misclassified cells and doublets using a first-pass SCDC run to improve robustness. Third, it allows single-subject scRNA-seq input, in which cross-subject variance cannot be directly estimated. (Refer to STAR Methods for more details.) However, since MDA-MB-468 and MCF-7 are both human breast cancer cell lines with relatively similar transcriptomic profiles, deconvolution of the bulk mixture by SCDC in a single run fails to estimate the correct relative proportions. To solve this issue, we applied the tree-guided deconvolution procedure proposed by MuSiC (Wang et al., 2019) to separate the closely related cell types. Refer to Supplemental Information for details.

The estimated cell-type proportions by SCDC with the tree-guided approach are 0.64:0.26:0.11, close to the ratio of 6:3:1 with a Pearson correlation of 0.991 (Figure 3B). We also benchmarked SCDC against Bseq-SC (Baron et al., 2016), CIBERSORTx (Newman et al., 2019), Bisque (Jew et al., 2019), and MuSiC (Wang et al., 2019), and showed that, even without ENSEMBLE, SCDC achieved the highest correlation coefficient. This is consistent with the simulations results shown in Table S1: overall, SCDC achieved the most accuracte deconvolution results when only one single-cell reference set was available.

### Performance on Real Dataset #2: Human Pancreatic Islets

To demonstrate the proposed ENSEMBLE framework when multiple reference datasets are available, we used SCDC to deconvolute 77 bulk RNA-seq samples of human pancreatic islets, of which 51 are from healthy individuals and 26 are from diabetic individuals (Fadista et al., 2014). Two scRNA-seq reference datasets were adopted, each harvesting six cell types of interest: alpha, beta, delta, gamma, acinar, and ductal cells (Baron et al., 2016, Åsa Segerstolpe et al., 2016). To allow the basis matrix **B** to reflect the potentially different gene expression patterns between the cases and controls, we performed the ENSEMBLE weight selection procedures separately for the samples from the two classes. The final ENSEMBLE weights for the two reference datasets were derived using both least absolute deviation (LAD) regression and grid search. Table 1 shows the final weights for the single-cell reference from Baron et al. (2016), which vary from 0.42 to 0.45 for the healthy samples and 0.49 to 0.52 for the diabetic samples. Figure 4A shows the cell-type proportions estimated with ENSEMBLE compared to the cell-type proportions estimated using single reference sets without ENSEMBLE. SCDC recovered to much higher levels the grossly underestimated fractions for beta cells by Baron et al. (2016), in concordance with the previous report by Cabrera et al. (2006). In addition, our results suggested that the beta cell proportions were slightly larger in the healthy donors than in the diabetic donors, although the difference was insignificant with a *p*-value of 0.1007.

**Table 1.**
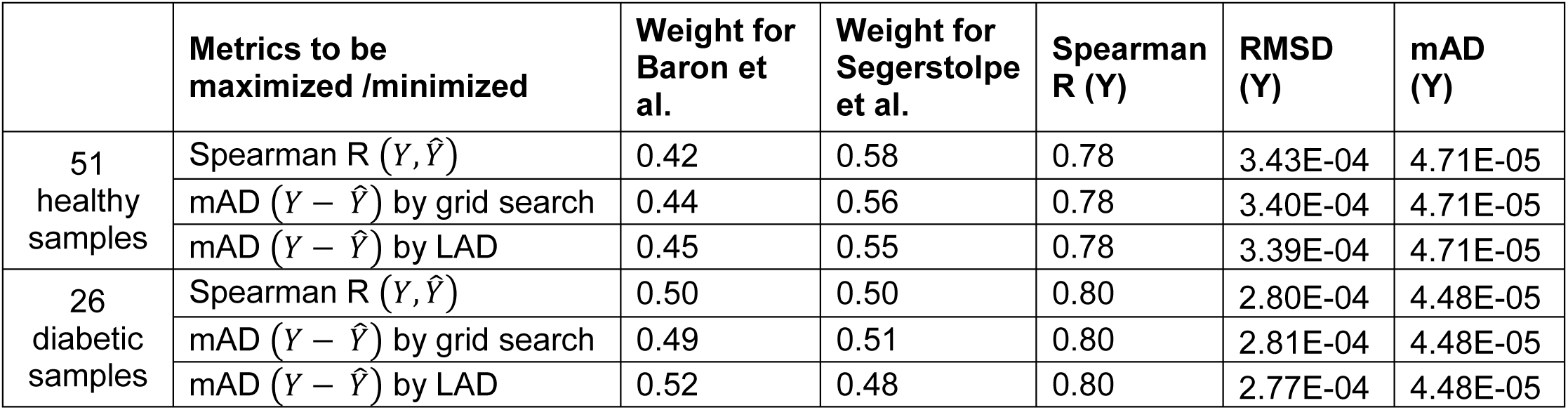
ENSEMBLE weight selection results for the human pancreatic islet bulk samples. The weights are presented separately for 51 healthy donors and 26 diabetic donors. SCDC selects weights that maximize the Spearman correlation of **Y** and or minimize the mAD of **Ŷ**, via grid search or least absolute deviation (LAD) regression.

**Figure 4.**
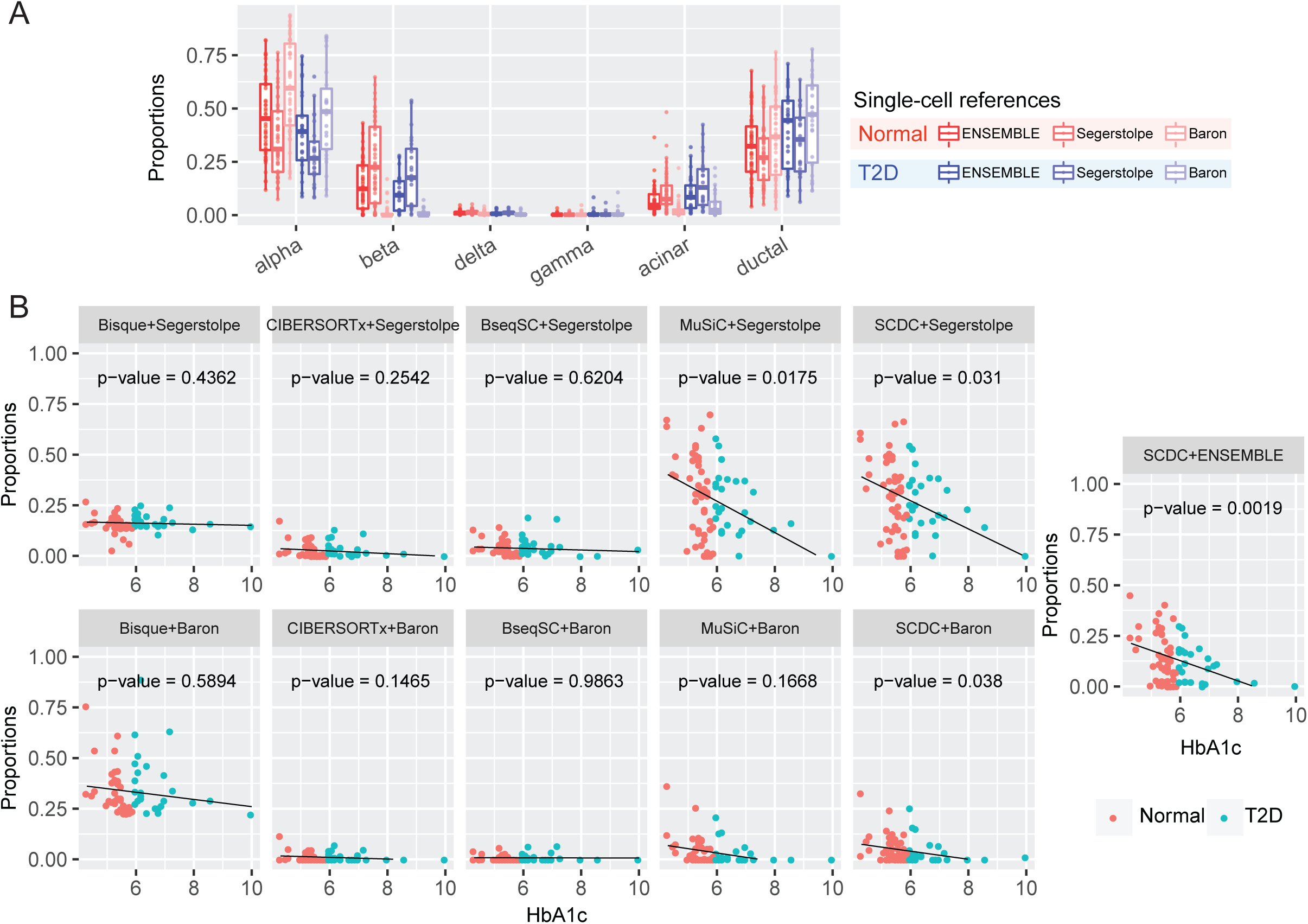
Gene expression deconvolution of human pancreatic islet samples. **A:** Estimated pancreatic islet cell-type composition in healthy and type 2 diabetic (T2D) human samples. The boxplot shows discrepancies in the deconvoluted proportions across different reference datasets. The ENSEMBLE method recovered the grossly underestimated beta cell proportions by deconvolution using only scRNA-seq data from Baron et al. (2016). **B:** Association of beta cell proportions and HbA1c levels by a linear model: beta cell proportion ∼ HbA1c + age + BMI + sex. Each benchmarked method was applied using reference datasets from Baron et al. (2016) and Åsa Segerstolpe et al. (2016) separately. The ENSEMBLE method by SCDC is additionally appiled using both reference datasets simultaneously. Bisque, CIBERSORTx, and BseqSC fail to recover the previously reported negative correlations; SCDC with ENSEMBLE returns the most significant *p*-value.

To evaluate the performance of SCDC and to compare against other existing methods, we sought to replicate previous findings on the negative correlation between the levels of hemoglobin A1c (HbA1c, an important biomarker for type 2 diabetes) and the beta cell functions (Kanat et al., 2011, Hou et al., 2016). We constructed a linear model using the estimated cell-type proportions as the response variable and the other covariates (age, gender, BMI, and HbA1c) as predictors. Overall, the ENSEMBLE method used with SCDC led to a p-value of 0.0019 for the negative relationship between the HbA1c levels and the beta cell proportions, more significant than the p-values of 0.031 and 0.038 from deconvolutions by SCDC without ENSEMBLE (Table S2, Figure 4B). Other existing methods – Bisque (Jew et al., 2019), CIBERSORTx (Newman et al., 2019) and BseqSC (Baron et al., 2016) – failed to recover the expected negative correlations, no matter which scRNA-seq reference dataset was adopted, and MuSiC (Wang et al., 2019) returned insignificant associations for the scRNA-seq reference dataset from Baron et al. (2016) (Figure 4B). In sum, the cell-type proportion estimates via ENSEMBLE more accurately reproduced the previously reported association between two orthogonal measurements.

### Performance on Real Dataset #3: Mouse Mammary Gland

We further illustrate the performance of SCDC on a dataset of mouse mammary gland. Figure 5A gives an overview of the experimental design. For this experiment, mouse mammary glands were harvested from two 12-week-old FVB/NJ mice, FVB3 and FVB4. Bulk RNA-seq was performed on the fresh frozen tissues. Meanwhile, single-cell suspension was prepared for the two samples; both scRNA-seq by 10X Genomics and bulk RNA-seq were performed on the pooled cell suspensions. (Refer to STAR Methods for details on experimental setup including animal model, cell suspension preparation, library preparation, and sequencing.) To illustrate the ENSEMBLE method for deconvolution, we adopted another single-cell reference dataset of mouse mammary glands from Tabula Muris (Consortium et al., 2018), generated by the microfluidic droplet-based method (see Key Resources Table). For clarity, the scRNA-seq data generated at the Perou Lab will be denoted as “Perou” and the scRNA-seq data from Tabula Muris will be denoted as “T. Muris”; the bulk RNA-seq data generated from the fresh frozen tissue will be denoted as “fresh frozen” and the bulk RNA-seq data from the pooled suspended cells will be denoted as “10X bulk.” We aimed to use SCDC to deconvolute each of the two bulk RNA-seq samples using the two scRNA-seq reference sets.

**Figure 5.**
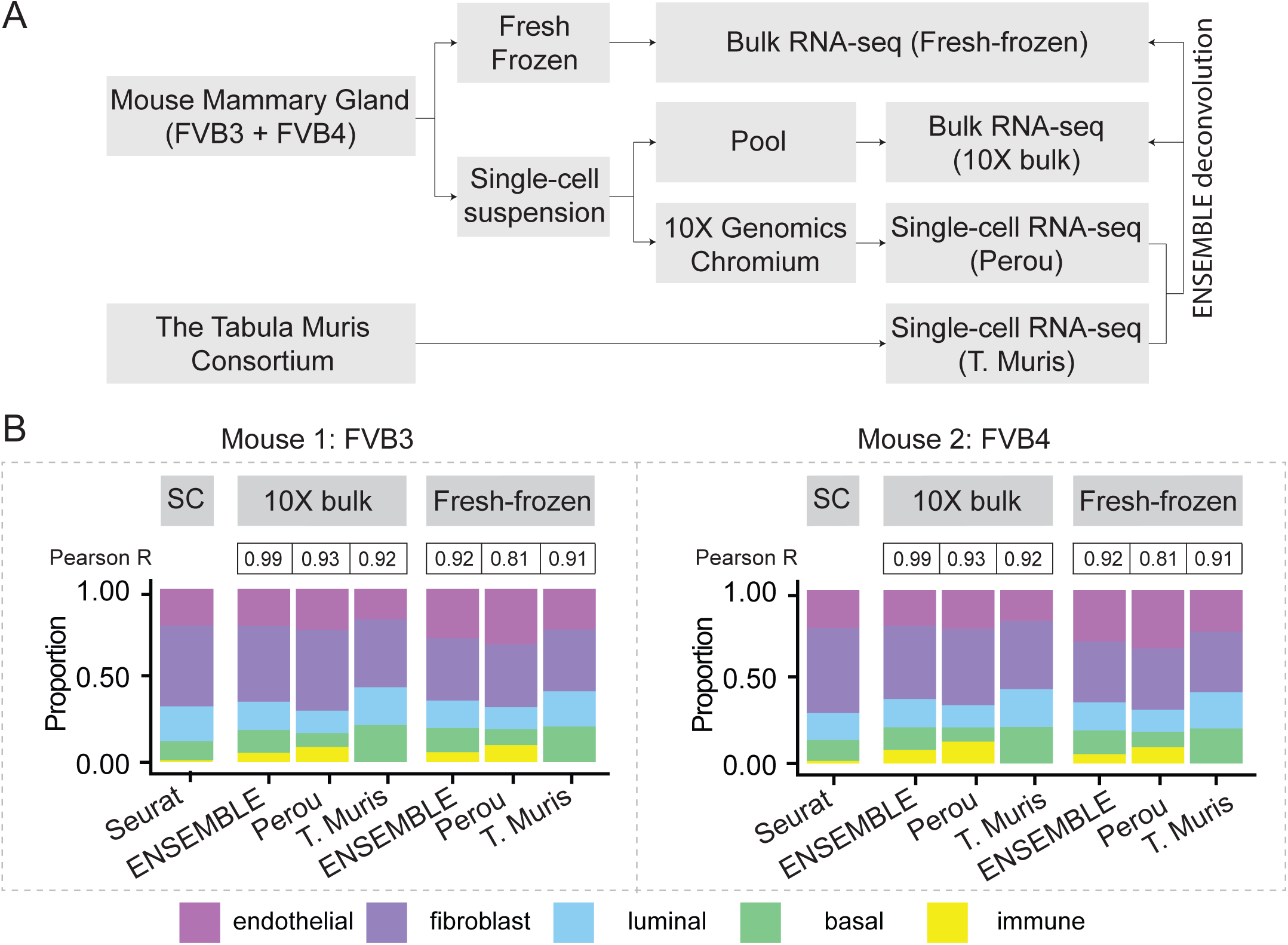
Gene expression deconvolution of mouse mammary gland samples. **A:** Flowchart of experimental design. Mouse mammary glands from two replicates, FVB3 and FVB4, were processed in two ways to generate both fresh-frozen bulk samples and single-cell suspensions. The single-cell suspensions were further divided into two parts, one for scRNA-seq by 10X Genomics, and the other for pooled bulk RNA-seq. To deconvolute the bulk samples through ENSEMBLE, another scRNA-seq dataset of mouse mammary gland from the Tabula Muris Consortium was adopted. **B:** Bulk gene expression deconvolution with and without ENSEMBLE. Pearson correlation of the cell-type proportions estimated by deconvolution and by scRNA-seq are shown. The ENSEMBLE method results in higher correlations for both bulk samples of the two replicates.

Following bioinformatic pre-processing (refer to STAR Methods for details), we first adopted Seurat (Butler et al., 2018) to perform single-cell clustering for the two scRNA-seq datasets, Perou and T. Muris, and then applied additional quality control (QC) procedures (outlined in the Methods section). The final cell types of interest consisted of immune, endothelial, fibroblast, luminal cells, and basal cells; t-SNE visualization is shown in Figure S3. As with the example of the three-cell-line mixture, we observed cell types with transcriptomic profiles that were highly similar (Figure S4A); we therefore adopted a tree-guided approach for deconvolution (Wang et al., 2019) in order to distinguish the closely related cell types (Figure S4B-C). This two-step deconvolution approach was applied using the Perou and T. Muris scRNA-seq references, respectively. Through ENSEMBLE, SCDC chose dataset-specific weights, which are shown in Table 2. As expected, a higher weight was assigned to the Perou reference dataset, which was from the same source as the bulk samples.

**Table 2.**
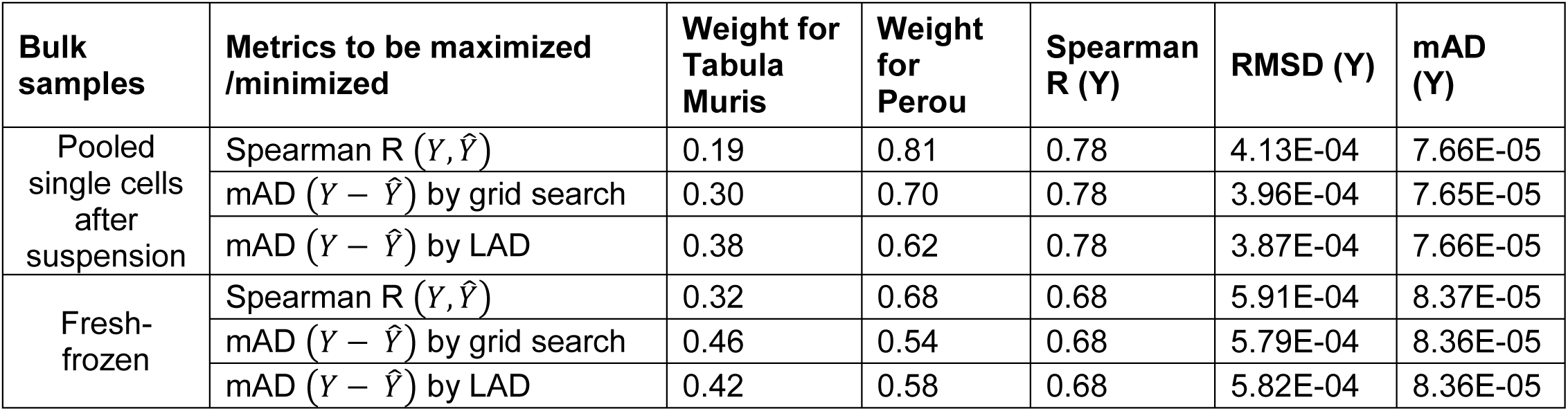
ENSEMBLE weight selection results for the mouse mammary gland bulk samples. The single-cell reference dataset from the same source as the bulk samples is more heavily weighted.

Figure 5B shows the final deconvolution results, both with and without ENSEMBLE, of the two bulk samples. The figure also includes Pearson correlations between the cell-type proportions estimated by scRNA-seq and those estimated by deconvolution. We found that the ENSEMBLE method produced higher correlation coefficients than approaches that use only one scRNA-seq dataset as reference (Figure 5B). This finding demonstrates the advantage of integrating data through SCDC. We also found that, compared to the fresh frozen bulk samples, the deconvoluted cell-type proportions from the 10X bulk samples were more highly correlated with the scRNA-seq fractions (Figure 5B). While the decrease of correlation coefficient from 0.99 to 0.92 is reassuring due to the order of the experiments, it also strikingly indicates a potential cell type-specific bias introduced by the 10X Genomics protocol, for it has been previously reported that adipocyte cells tend to get lost during the single-cell library preparation step (Kessenbrock et al., 2018). As such, cell-type proportions from the single cell experiment do not necessarily reflect those in the bulk tissues due to the sampling bias and the technical artifacts that are associated with the library preparation and sequencing step of scRNA-seq (Hwang et al., 2018). This makes *in silico* deconvolution a compelling approach to unbiased recovery of true underlying cell-type composition.

## Discussion

Here, we propose a method for deconvoluting bulk RNA-seq data accurately by exploiting multiple scRNA-seq reference datasets through ENSEMBLE. We show that such data integration leads to higher deconvolution accuracy via both extensive simulations and experimental validations. Existing batch correction methods for scRNA-seq data either do not return a gene expression matrix that is adjusted for batch effect (Butler et al., 2018) or return one with a drastically different range of measurements (Haghverdi et al., 2018). These drawbacks make them insufficient and infeasible for joint deconvolution analysis. SCDC does not directly address this nontrivial issue to correct for batch effect; rather, it opts to integrate results from all scRNA-seq datasets with different weights, so as to reflect the degree of similarity between the bulk data and the reference data. The ultimate goal is to return a deconvolution result as close to the truth as possible. Similarly, for bulk RNA-seq data, which can also potentially harbor batch effects, SCDC can select an optimal combination of scRNA-seq reference sets for each sample separately to achieve more accurate cell-type decomposition. In addition, while some methods may require paired bulk-tissue and single-cell RNA-seq data from the same individuals (Jew et al., 2019), SCDC has no such requirement due to its robustness to technical variability.

While in this paper we have focused on integrating results from multiple scRNA-seq data sets, the same framework can be applied to integrate results from different deconvolution methods. In Table S1, we showed that no one method universally performed better than the others across all simulation setups. To address this instability issue, SCDC’s weighting principle can be applied similarly, where different weights are assigned to different deconvolution methods.

Identifying cell-type composition of disease-relevant tissues allows identification of cellular targets for treatment and offers a better understanding of disease mechanism. For downstream analysis following deconvolution, hypothesis testing on differential gene expression in a case-control setting needs to account for the variability of cell-type composition. As Shen-Orr et al. (2010) have described, differential gene expression analysis in the presence of cellular heterogeneity can be performed through the following testing schemes: (i) whole tissue differences (i.e., testing on **Y);** (ii) differences in cell-type compositions (i.e., testing on **P);** (iii) differences in cell type-specific gene expression patterns (i.e., testing on **B**_:*k*_ for each cell type *k*); (iv) differences in cell type-specific gene expression patterns while accounting for cell-type proportions (i.e., testing on **B**_:*k*_**P**_*k*:_ for each cell type *k*); and (v) an omnibus test across all cell types (i.e., testing on **B** across all cell types simultaneously). All of these testing schemes (except for the testing on **Y** by traditional methods developed for bulk RNA-seq data) must be adapted when scRNA-seq data is used to aid deconvolution: neither **B** nor **P** is pre-known, and one must take into consideration their estimation uncertainties through deconvolution. The questions of how to jointly perform differential testing when multiple scRNA-seq datasets are available and how to jointly model both bulk and single-cell RNA-seq data (Zhu et al., 2018) with high computational efficiency require further investigation.

## Acknowledgments

This work was supported by the National Institutes of Health (NIH) grant T32 ES007018 (to EU), R01 HL129132 (to YL), R01 GM105785 (to FZ), P30 ES010126 (to FZ), P01 CA142538 (to YJ), R35 GM118102 (to YJ), UL1 TR002489 (to YJ), National Cancer Institute (NCI) Breast SPORE program P50 CA5822 (to CMP), R01 CA148761 (to CMP), Breast Cancer Research Foundation (to CMP), a developmental award from the UNC Lineberger Comprehensive Cancer Center 2017T109 (to YJ), and a pilot award from the UNC Computational Medicine Program (to YJ).

## Author Contributions

FZ and YJ initiated and envisioned the study. MD, FZ, and YJ formulated the model. MD implemented the algorithm and performed simulation studies. AT and CMP envisioned and performed the cell line mixing and normal mammary gland experiments. All authors performed real data analysis. MD and YJ wrote the manuscript, which was further edited and approved by all authors.

## STAR Methods

### Key Resources Table

Separately attached.

### Contact for Reagent and Resource Sharing

Further information and requests for resources and reagents should be directed to and will be fulfilled by Charles M. Perou, Fei Zou, and Yuchao Jiang

### Experimental Model and Subject Details

#### Cell-line mixture

MCF-7 and MDA-MB-468 cells were purchased from ATCC. Human dermal fibroblasts were isolated from skin. All cell lines were maintained independently in culture medium DMEM (Gibco) supplemented with 10% FBS (Millipore) and 1% penicillin-streptomycin (Gibco) and grown in incubators maintained at 37 °C with 5% CO_2_. Cells were mixed together so that MCF-7 cells comprised 60% of the mixture, MDA-MB-468 cells comprised 30% of the mixture, and dermal fibroblasts comprised 10% of the mixture.

#### Animal model

All animal studies were performed with approval and in accordance with the guidelines of the Institutional Animal Care and Use Committee (IACUC) at the University of North Carolina at Chapel Hill. Female FVB/NJ mice were obtained in collaboration with the UNC Lineberger Comprehensive Cancer Center (LCCC) Mouse Phase I Unit (MP1U). Animals were cared for according to the recommendations of the Panel on Euthanasia of the American Veterinary Medical Association. Mice were housed in a climate controlled Department of Laboratory Animal Medicine facility with a 12 h light:dark cycle and ad libitum access to food and water (Qin et al., 2016). The mammary glands were harvested at 12 weeks for FVB/NJ mice.

### Method Details

#### Cell suspension preparation

The FVB/NJ mammary glands were placed in 10 ml of a digestion medium containing EpiCult™-B Mouse Medium Kit (#05610, StemCell Technologies), Collagenase/Hyaluronidase (#07912, StemCell Technologies), and 1% penicillin-streptomycin (Gibco). The mammary gland was digested overnight in a thermocycler maintained at 37 °C with continuous rotation. The cell pellets retrieved from these suspensions were treated with a 1:4 solution of hanks balanced salt solution (HBSS) and ammonium chloride to remove the RBCs. After RBC removal, the cell suspensions were trypsinized with 0.05% Trypsin and a mix of Dispase and DNAse. A portion of this cell suspension was stained with trypan blue and counted using the Countess Automated Cell Counter (lnvitrogen). Based on the counting, the cells were diluted to the appropriate cell stock concentration for running on the 10X Chromium machine. Based on the 10X Genomics pre-defined cell stock concentrations, each experiment was run to retrieve ∼5000 cells after the single-cell experiment. The remaining cell stock solution was used for making bulk mRNA seq libraries.

#### Single-cell library construction, sequencing, and bioinformatics pipeline

The cell suspensions were loaded on a 10X Genomics Chromium instrument to generate single-cell gel beads in emulsion (GEMs) for targeted retrieval of approximately 5000 cells. scRNA-Seq libraries were prepared following the Single Cell 3’ Reagent Kits v2 User Guide (Manual Part # CG00052 Rev A) using the following Single Cell 3’ Reagent Kits v2: Chromium™Single Cell 3’ Library & Gel Bead Kit v2 PN-120237, Single Cell 3’ Chip Kit v2 PN-120236, and i7 Multiplex Kit PN-120262” (10X Genomics). Libraries were run on an Illumina HiSeq 4000 as 2 × 150 paired-end reads. The Cell Ranger Single Cell Software Suite (version 1.3) was used to perform sample de-multiplexing, barcode and unique molecular identifiers (UMI) processing, and single-cell 3’ gene counting. All scRNAseq data by 10X Genomics are available at GEO database (GSE136148).

#### Bulk mRNA-seq pre-processing

RNA was isolated using the RNeasy Mini Kit (#74104, Qiagen) according to manufacturer protocol. mRNA quality was assessed using the Agilent Bioanalyzer and libraries for mRNA-seq were made using total RNA and the lllumina TruSeq mRNA sample preparation kit. Paired end (2×50bp) sequencing was performed on the Illumina HiSeq 2000/2500 sequencer at the UNC High Throughput Sequencing Facility (HTSF). Resulting fastq files were aligned to the mouse mm10 reference genome using the STAR aligner algorithm (Dobin et al., 2013). Resulting BAM files were sorted and indexed using Samtools (Li et al., 2009) and QC was performed using Picard *(Picard*, 2019). Transcript read counts were determined using Salmon (Patro et al., 2015). Genes with zero read counts across all samples were removed. All bulk mRNAseq data is available at GEO database (GSE136148).

#### Clustering quality control of scRNA-seq data

To construct the basis matrix B from the single-cell reference dataset, SCDC takes as input gene expression measurements and cluster memberships of all cells that are sequenced by scRNA-seq. While much efforts have been devoted to cell type clustering by scRNA-seq, it has been shown that different approaches can potentially generate varying single-cell cluster assignments (Huh et al., 2019). To make SCDC robust to single-cell clustering, a quality control procedure is performed as a first step to remove cells with questionable cell-type assignments, as well as cells with low library preparation and sequencing quality. Specifically, each single cell is treated as a “bulk” sample and its cell-type composition can be derived by a first-pass run of SCDC. For well classified cells with good quality, the estimated proportions should be sparse and contain a single non-zero estimate close to one; for questionable cells such as doublets, the estimated proportions would not result in a unique cluster assignment (Figure S5A). Therefore, we remove cells whose estimated cell-type proportions have a maximum less than a user-defined threshold (Figure S5B). After this initial QC step of the single-cell input, the Pearson correlation of the actual and the deconvoluted cell-type proportions is improved for simulation runs, especially when pseudo-bulk samples and reference datasets are from different sources (Table S1).

#### Construction of basis matrix differs from MuSiC

For deconvolution using each single-cell reference dataset, SCDC estimates cell-type proportions following the W-NNLS framework proposed by MuSiC (Wang et al., 2019), but differs in the way of calculating the basis matrix. The contribution of each subject to the construction of a basis matrix may vary according to the data quality (Figure S6). Hence, maximal variance weight (MVW) per gene is calculated to reflect the data quality (Wilson et al., 2018). In detail, using scRNA-seq data, SCDC first estimates 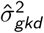 which captures the cross-cell variation for gene *g* of cell type *k* within individual *d.* Within-subject variance for subject dis then calculated as 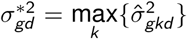 and the maximal variance weight Δ_*gd*_ is given by:

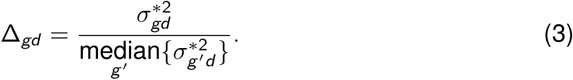

SCDC proceeds to scale the raw single-cell read count matrix by 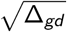 Under this construction, genes with larger variance will have larger variance weights. Larger variance weights ensure that residuals from such genes will have smaller impact on estimation of cell-type composition (Wilson et al., 2018). To control for excessively large or small variance weights, we set the bottom 15% of variance weights to be the 15th percentile variance weight, and similarly, the top 15% of variance weights are replaced by the 85th percentile variance weight. The rest of the estimation procedure largely follows MuSiC. The performances of SCDC and MuSiC were compared via simulations by Pearson correlation, RMSD and mAD between 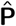 and **P** shown in Table S1.

#### ENSEMBLE: a linear combination of deconvolution results

Assume *R* single-cell reference datasets are available for the tissue of interest. For each reference dataset *r* ∈ {1, …, *R*}, SCDC deconvolutes the bulk gene expression data as a matrix decomposition problem. Let **P**_*r*_ and **B**_*r*_ denote the cell-type proportion matrix and the basis matrix using the rth reference dataset, respectively. The bulk gene expression **Y** can be deconvoluted into the form of **Y** = **B**_*r*_**P**_*r*_ + *ϵ*_*r*_ with a reference-specific error term *ϵ*_*r*_. The predicted gene expression levels from the *r*^*th*^ reference dataset is 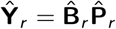 In the ENSEMBLE step, SCDC aims to solve for equation (2). As we stated in the Result session, we make the assumption that the solutions for equation (1) and (2) are approximately equivalent based on the concordance between the metrics on the cell-type proportions (Pearson correlation and mAD between 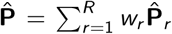 and **P)** and the metrics on the gene expression levels (Spearman correlation, RMSD, and mAD between 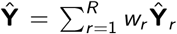 and **Y** via simulations (Figure 2, Figure S2). See Supplemental Information for equation details. In practice, SCDC, by default, chooses the L1 norm of **(Y** − **Ŷ)** as the criteria for ENSEMBLE weight selection.

For optimization, we can redirect the problem to least absolute deviations (LAD) regression with constraints on the weights (w_1_, …, w_*R*_):

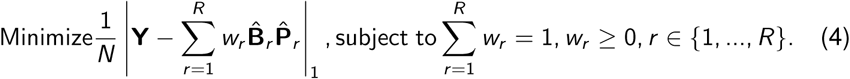

LAD regression does not have an analytical solving method (Vanderbei, 2001), hence we applied the method adopted by Osorio et al. (2017). While solving for *w*_*r*_’s, an LAD regression with no constraint is first fit. Any negative *ŵ*_*r*_ is set to zero, and the estimates are finally scaled to satisfy the constraint. Since the rescaling step can be problematic, SCDC additionally adopts another numerical method via grid search to determine the optimal ENSEMBLE weights. However, the grid search method might be computationally inefficient if more than three reference datasets are used and the search step size is set too small. Regardless, the optimal weights selected by LAD and by grid search generally agree with each other, as we showed in real data analysis (Table 1, Table 2).

### Data and Software Availability

SCDC is compiled as an open-source R package available at https://github.com/meichendong/SCDC, together with vignettes and toy examples for demonstration.

## Supplemental Information

### Evaluation Measurement

The metrics we used for method evaluation include root-mean-square deviation (RMSD), mean absolute deviation (mAD), Pearson correlation, and Spearman correlation. Given a parameter *z* and its estimator 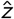, these metrics can be defined as:

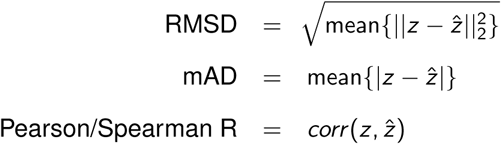

### Quality Control and Clustering of scRNA-seq Data

For single cells from the three cell-line experiment, cells with a high percentage of mitochondrial gene expressions were filtered out. Genes with lengths greater than 200kb, ribosomal genes, and genes with undetectable expressions were filtered out. Seurat was applied for single cell clustering: genes detected in at least three cells were kept; cells with less than 200 genes detected were filtered out; the number of genes detected and the number of UMIs were regressed out in the scaling procedure; ‘FindClusters’ was applied using the first twenty principal components, with resolution parameter set from 0.6 to 1. Finally, cell types were annotated according to previously reported marker genes.

For the mouse mammary gland data, single cell clustering was performed within each subject separately. In addition to the Seurat clustering procedures described above, the percentage of cell-cycle gene expressions was also regressed out when scaling the gene expression matrix. Epithelial cells were first identified as a major cluster and were further subgrouped into luminal and basal cells. ‘FindMarkers’ function was applied to each pair of cell types, and the number of marker genes from each pair was used to determine whether or not to combine the two clusters.

### Two-Level Deconvolution

Similar to MuSiC (Wang et al., 2019), for cases where closely related cell types are present in the data, SCDC adopts a two-step approach, which first separates remotely connected cell types and, in the second step, dissociates cell types that share high similarities. However, there is no consensus on how to determine the order of deconvolution, especially when multiple scRNA-seq datasets are available. To solve this, we employ **MNN** (Haghverdi et al., 2018) to correct for batch effect and to calculate a basis matrix from the adjusted data. Hierarchical clustering is applied to determine the relationship between the cell types of interest. The hierarchical structure is further used to guide the two-step approach for deconvolution. For the mouse mammary gland dataset, the first-round deconvolution separates cluster 1 = {immune cells} from cluster 2 ={endothelial, fibroblast, basal, luminal cells} and the second-round deconvolution further separates the cell types in cluster 2 (Figure S4A). Within each level of deconvolution, differentially expressed genes are first identified by Wilcoxon rank-sum test with multiple testing correction and then used as input.

### Deconvolution Using Single Cells from One Subject

To accommodate experimental designs using single cells from only one subject, we adapt the W-NNLS framework to calculate the gene-specific weights by within-subject variation only. Denote the cell-type proportion vector for bulk sample *d* as **P**_*d*_= (***P***_1*d*_, ***P***_*2d*_, …, ***P***_*Kd*_)^*T*^ and the normalized bulk gene expression as ***Y***_*d*_*= (Y*_1*d*_, *Y*_2*d*_, …, *Y*_G*d*_) ^*T*^ The gene-specific expression can be formalized as

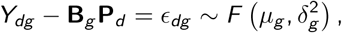

where **B**_*g*_ is the *g* ^*th*^ row in the basis matrix **B;** the residual term *ϵ*_*dg*_ follows a certain distribution *F* with mean *µ*_*g*_ and variance 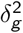;. Adjusting for the variance of residuals, we derive:

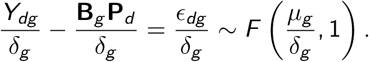

We can iteratively estimate the proportion vector **P**_*d*_ and derive the residual vector in the meantime. If two consecutive estimated proportion vectors 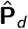 and 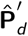 are equal, then we derive a consistent estimation result. That is, if 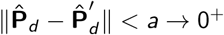 and 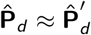 then

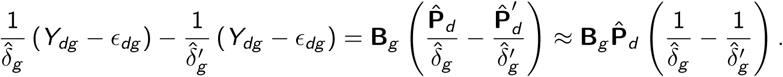

Hence, as the proportion estimates converge, we derive a final deconvolution result:

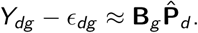

## KEY RESOURCES TABLE

| REAGENT or RESOURCE | SOURCE | IDENTIFIER |
| --- | --- | --- |
| **Chemicals, Peptides, and Recombinant Proteins** |  |  |
| EpiCult™-B Mouse Medium Kit | StemCell Technologies | EpiCult™-B Mouse Medium Kit #05610 |
| Dispase (5 U/mL) | StemCell Technologies | Dispase (5 U/mL) #07913 |
| Ammonium Chloride Solution | StemCell Technologies | Ammonium Chloride Solution #07850 |
| Gentle Collagenase/Hyaluronidase | StemCell Technologies | Gentle Collagenase/Hyaluronidase # 07919 |
| DNase I Solution (1 mg/mL) | StemCell Technologies | DNase I Solution (1 mg/mL) # 07900 |
| **Critical Commercial Assays** |  |  |
| TruSeq Stranded mRNA | Illumina | TruSeq® Stranded mRNA Library Prep (48 Samples)<br>20020594 |
| Chromium Single Cell 3' Reagent Kits | 10x Genomics | Chromium Single Cell 3' GEM, Library & Gel Bead Kit v3, 4 rxns PN-1000092<br>Chromium Single Cell B Chip Kit, 48 rxns PN-1000073<br>Chromium i7 Multiplex Kit, 96 rxns PN-120262 |
| MACS Tumor Dissociation Kit, mouse | Miltenyi Biotec | Tumor dissociation kit # 130-096-730 |
| **Deposited Data** |  |  |
| ScRNA-seq of human pancreatic islet | Baron et al., 2016 | GEO: GSE84133 |
| ScRNA-seq of human pancreatic islet | Xin et al., 2016 | GEO: GSE81608 |
| ScRNA-seq of human pancreatic islet | Segerstolpe et al., 2016 | GEO: E-MTAB-5061 |
| Bulk RNA-seq of human pancreatic islet | Fadista et al., 2014 | GEO: GSE50244 |
| ScRNA-seq of mouse mammary gland | Tabula Muris, 2018 | GEO: GSE106273 |
| ScRNA-seq of mouse mammary gland | This paper | GEO: GSE136148 |
| Bulk RNA-seq of mouse mammary gland | This paper | GEO: GSE136148 |
| ScRNA-seq of three-cell-line mixture | This paper | GEO: GSE136148 |
| Bulk RNA-seq of three-cell-line mixture | This paper | GEO: GSE136148 |
| **Experimental Models: Cell Lines** |  |  |
| MCF7 | ATCC | MCF7 (ATCC® HTB-22™)<br>https://www.atcc.org/products/all/HTB-22.aspx |
| MDA-MB-468 | ATCC | MDA-MB-468 (ATCC® HTB-132™)<br>https://www.atcc.org/products/all/HTB-132.aspx |
| Human dermal Fibroblasts | Perou Lab | N/A |
| **Experimental Models: Organisms/Strains** |  |  |
| Mouse: FVB/NJ | The Jackson Laboratory | JAX: 001800 |
| Mouse: FVB-Tg(C3-1-TAg)cJeg/JegJ | The Jackson Laboratory | JAX: 013591 |
| **Software and Algorithms** |  |  |
| Seurat | Satija et al., 2015<br>Butler et al., 2018 | https://github.com/satijalab/seurat |
| MuSiC | Wang et al., 2019 | https://github.com/xuranw/MuSiC |
| Bseq-SC | Baron et al., 2016 | https://github.com/shenorrLab/bseqsc |
| Bisque | Jew et al., 2019 | https://github.com/cozygene/bisque |
| CIBERSORTx | Newman et al., 2019 | https://cibersortx.stanford.edu |
| SCDC | This paper | https://github.com/meichendong/SCDC |

## Supplemental Figure Titles and Legends

**Figure S1.**
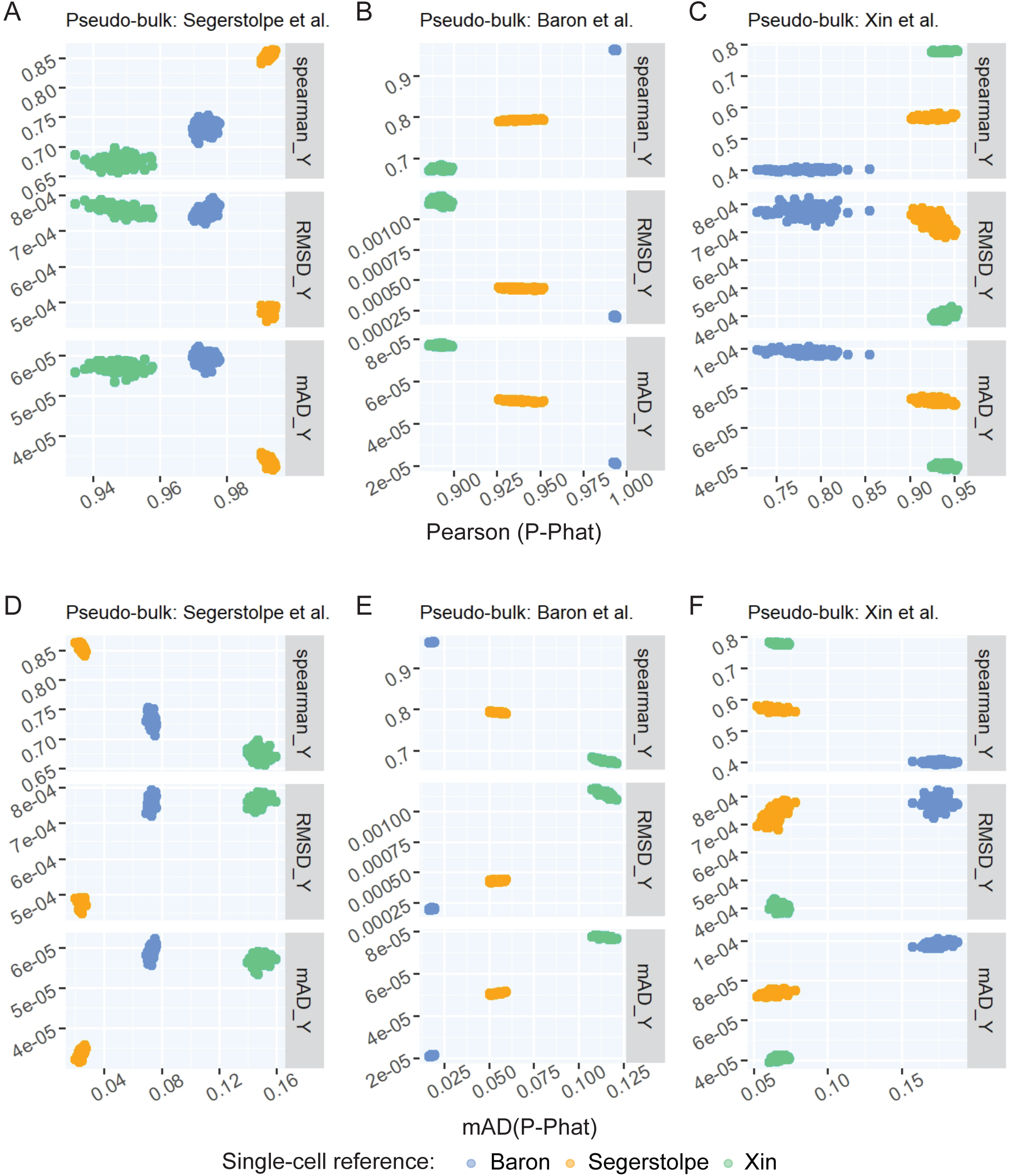
Empirical results via simulations show that the metrics on gene expression levels **Y** are good proxies for the metrics on cell-type proportions **P. A-C:** Prediction errors ‖**Y − Ŷ** ‖1 against Pearson correlation between cell-type proportions **P** and 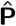 for pseudo-bulk samples constructed using single cells from Asa Segerstolpe et al. (2016), Baron et al. (2016), and Xin et al. (2016), respectively. **D-F:** Prediction errors ‖**Y − Ŷ** ‖1 against 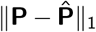 for pseudo-bulk samples constructed from Asa Segerstolpe et al. (2016), Baron et al. (2016), and Xin et al. (2016), respectively.

**Figure S2.**
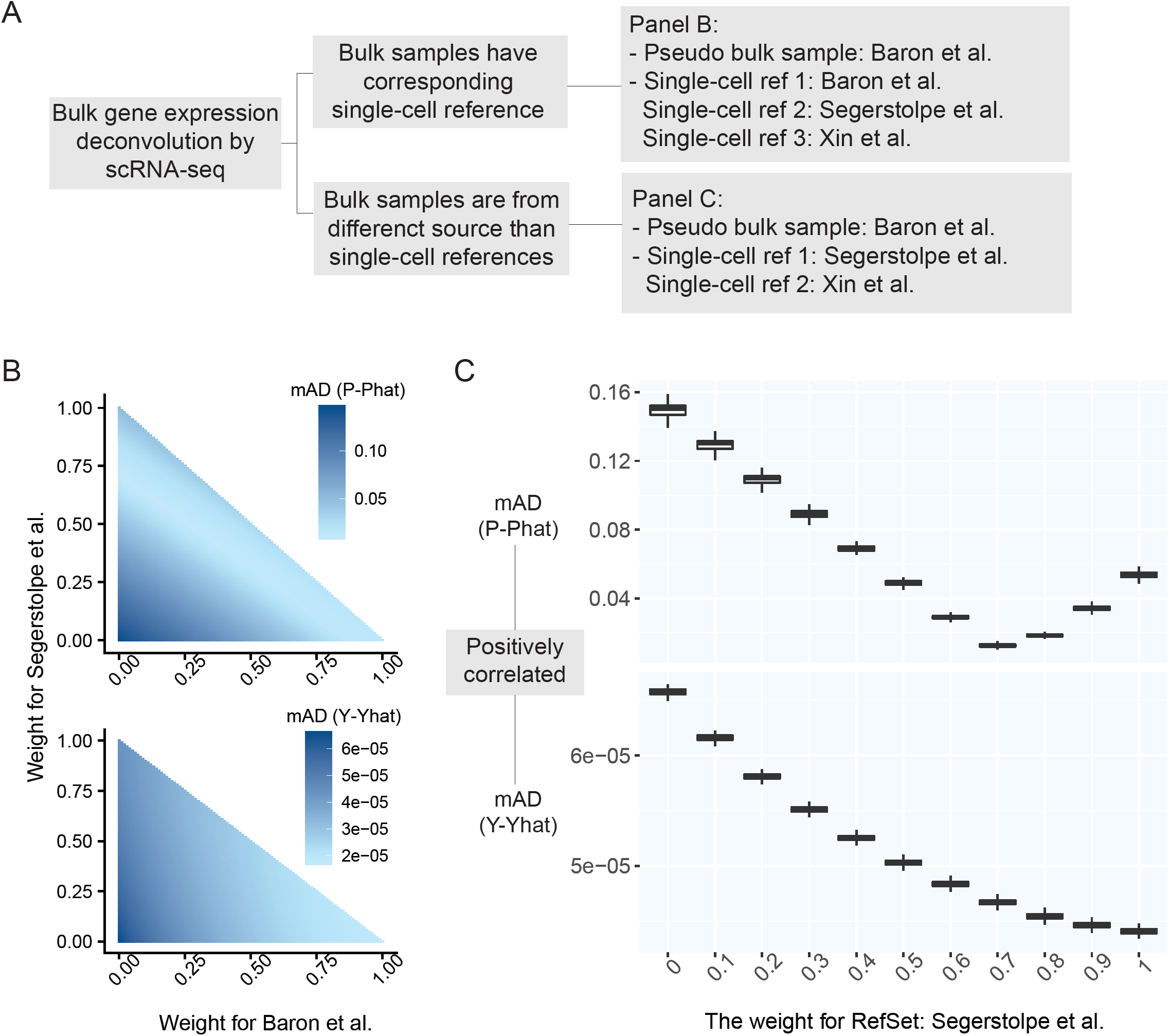
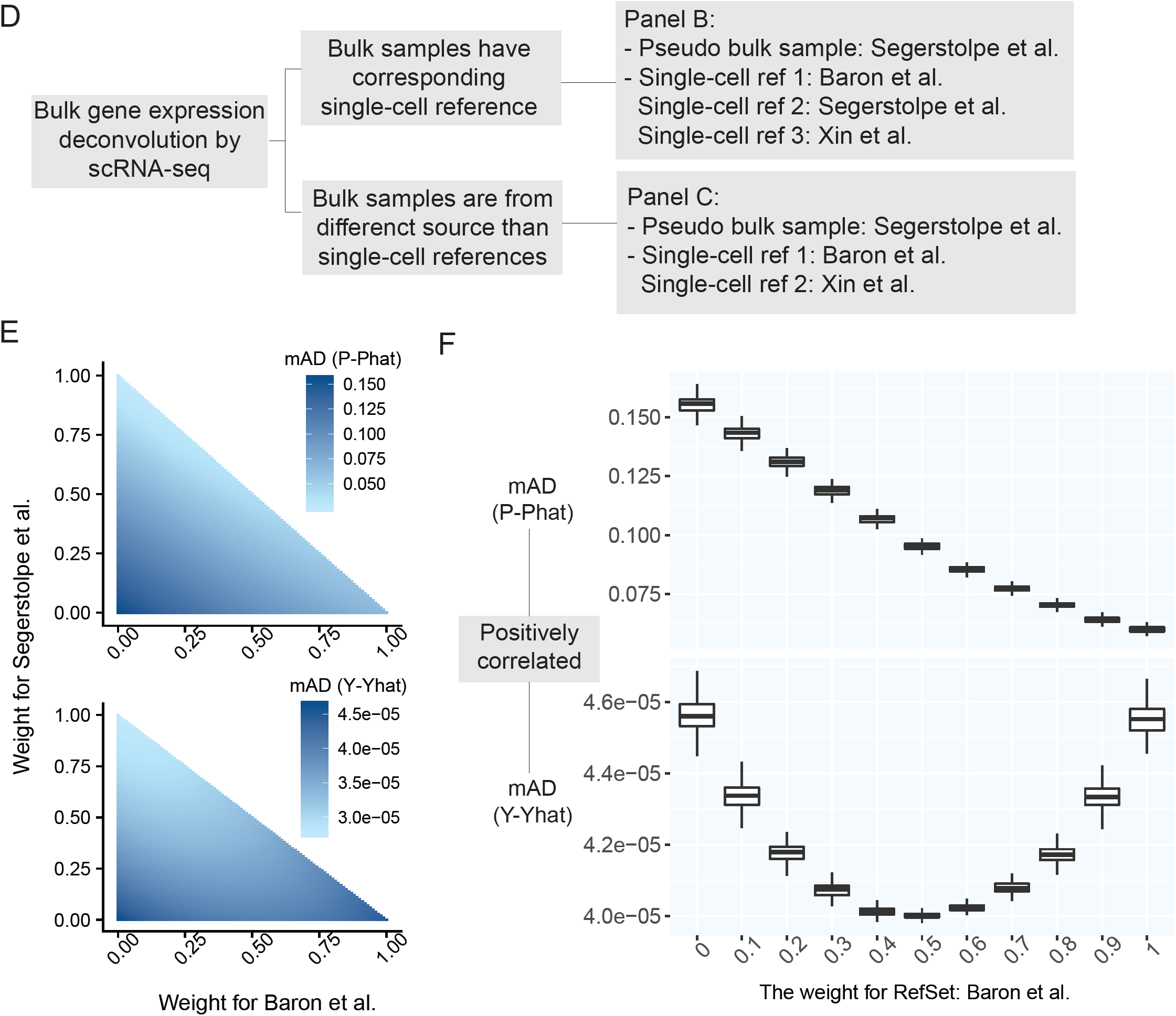
Prediction errors of **Y** serve as a surrogate for the estimation errors of P. The simulation setups differ from those in Figure 2. **A:** Outline of simulation setup, where single cells of human pancreatic islets from Baron et al. (2016) are aggregated to generate pseudo-bulk samples, whose cell-type pro-portions are known. We examine the results of deconvolution via ENSEMBLE under two settings, both with and without paired single-cell reference datasets. 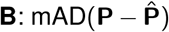 and mAD(**Y** − **Ŷ)** with three varying dataset-specific weights for deconvolution of bulk samples with paired scRNA-seq. The two metrics agreed on the assignment of the optimal weights: around (*ŵ*_1_, *ŵ*_2_, *ŵ*_3_) = (1,0,0) 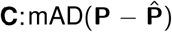 and mAD(**Y** − **Ŷ)** with two varying dataset-specific weights for deconvolution of bulk samples without paired scRNA-seq. The two metrics are highly correlated with varying weights for the reference dataset from Åsa Segerstolpe et al. (2016). **D:** Outline of simulation setup, where single cells of human pancreatic islets from Åsa Segerstolpe et al. (2016) are aggregated to generate pseudo-bulk samples, whose cell-type proportions are known. 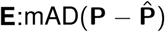 and mAD(**Y** − **Ŷ)** with three varying dataset-specific weights for deconvolution of bulk samples with paired scRNA-seq. The two metrics agreed on the assignment of the optimal weights to be around (*ŵ*_1_, *ŵ*_2_, *ŵ*_3_) = (0,1,0). 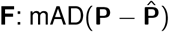 and mAD(**Y** − **Ŷ)** with two varying dataset-specific weights for deconvolution of bulk samples without paired scRNA-seq. While the two metrics do not share the same trend with the varying weights, the weight selected by mAD(**Y** − **Ŷ)** would achieve a mAD 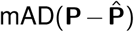 close to its smallest value.

**Figure S3.**
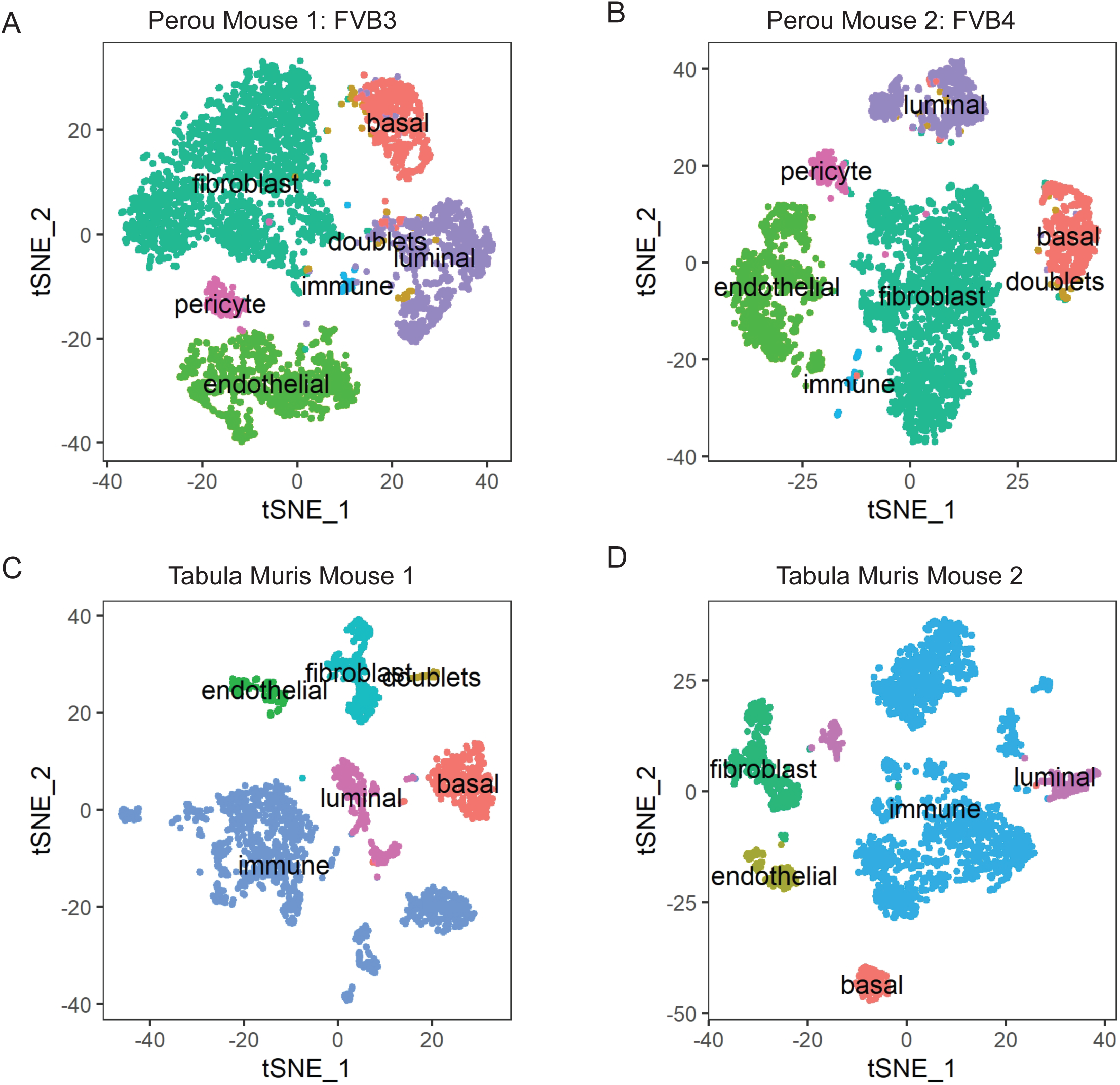
Single-cell clustering visualization by t-SNE. **A-B:** scRNA-seq data from the Perou Lab. **C-D:** scRNA-seq data from the Tabula Muris Consortium.

**Figure S4.**
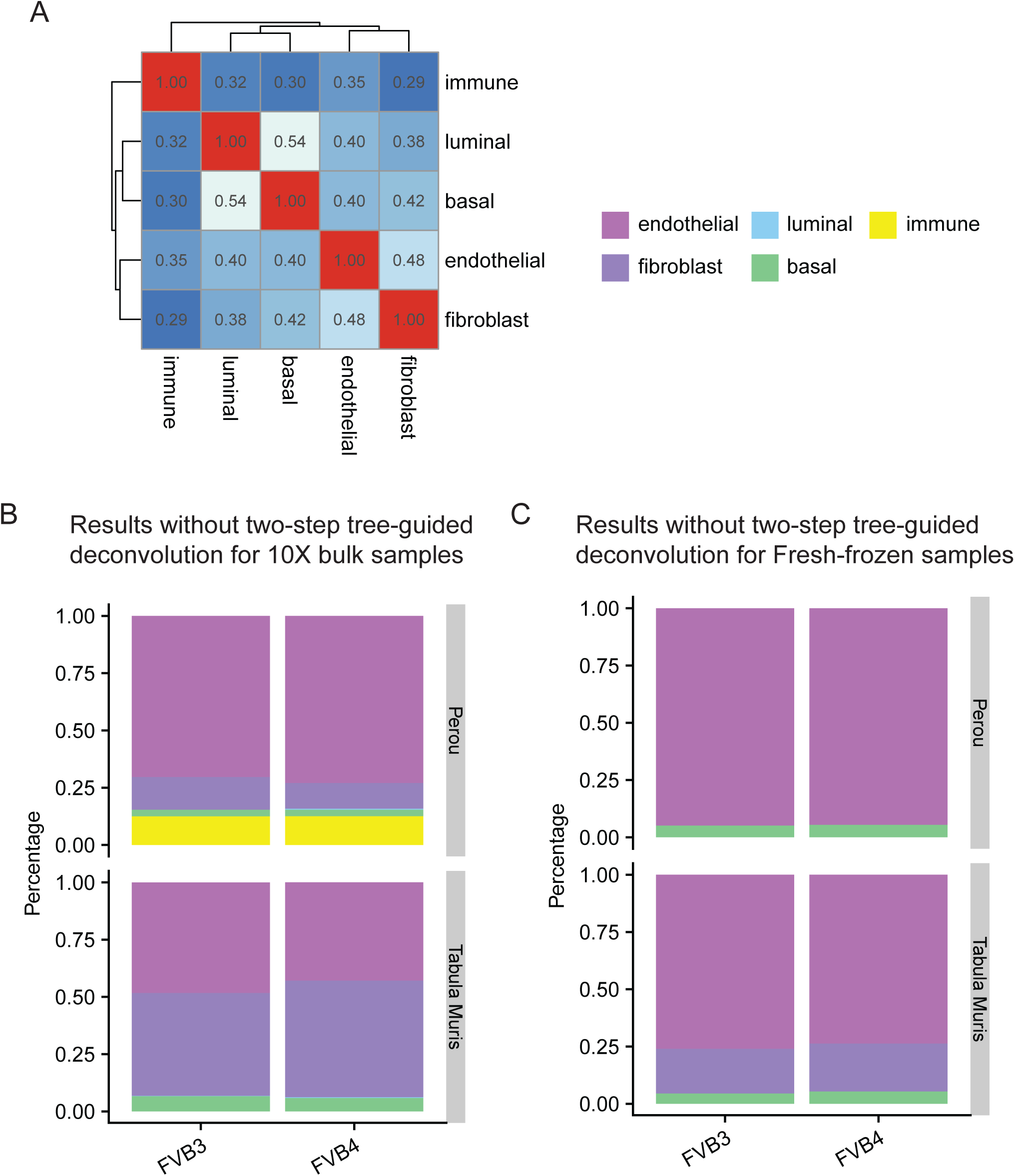
Deconvolution results without the tree-guided approach hardly separate closely related cell types. **A:** Pairwise correlation of cell-type-specific gene expression profiles estimated by scRNA-seq. **B:** Estimated cell-type proportions of mouse mammary gland 10X bulk samples without tree-guided approach. **C:** Estimated cell-type proportions of mouse mammary gland fresh-frozen bulk samples without tree-guided approach.

**Figure S5.**
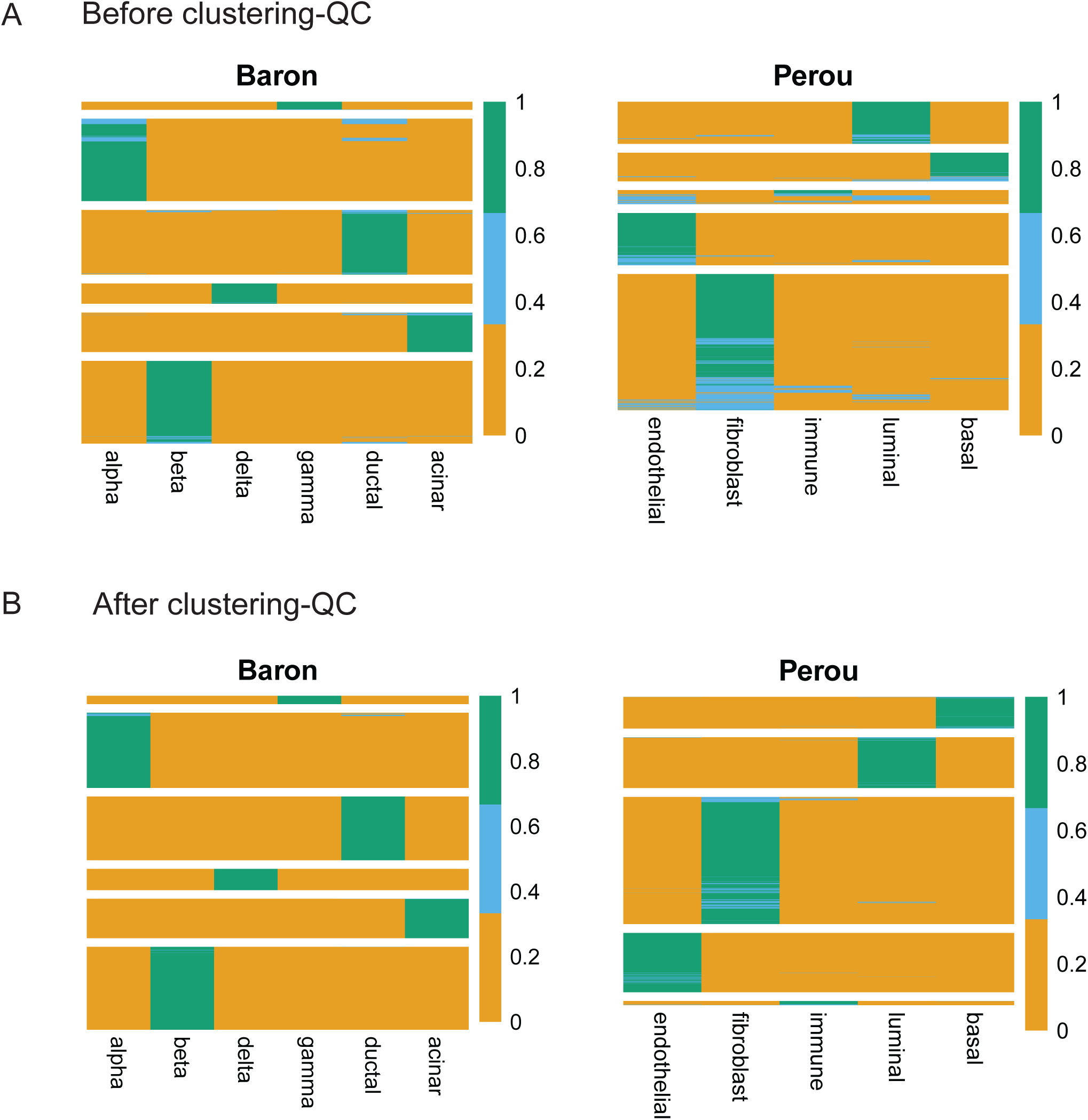
A first-pass SCDC run on the single-cell reference dataset removes potentially mislabeled cells and doublets. Each single cell is treated as a “bulk” sample and used as input for SCDC. The highly binary cell-type proportions indicate good data quality and reliable cell type clustering. Cells whose estimated cell-type proportions have a maximum less than a user-defined threshold (0.7 by default) are filtered out. These cells are potentially doublets, mis-classified, poorly sequenced, or from cell types not of interest. **A:** A first-pass SCDC run using cells as “bulk” samples. **B:** Unique cell identities after QC.

**Figure S6.**
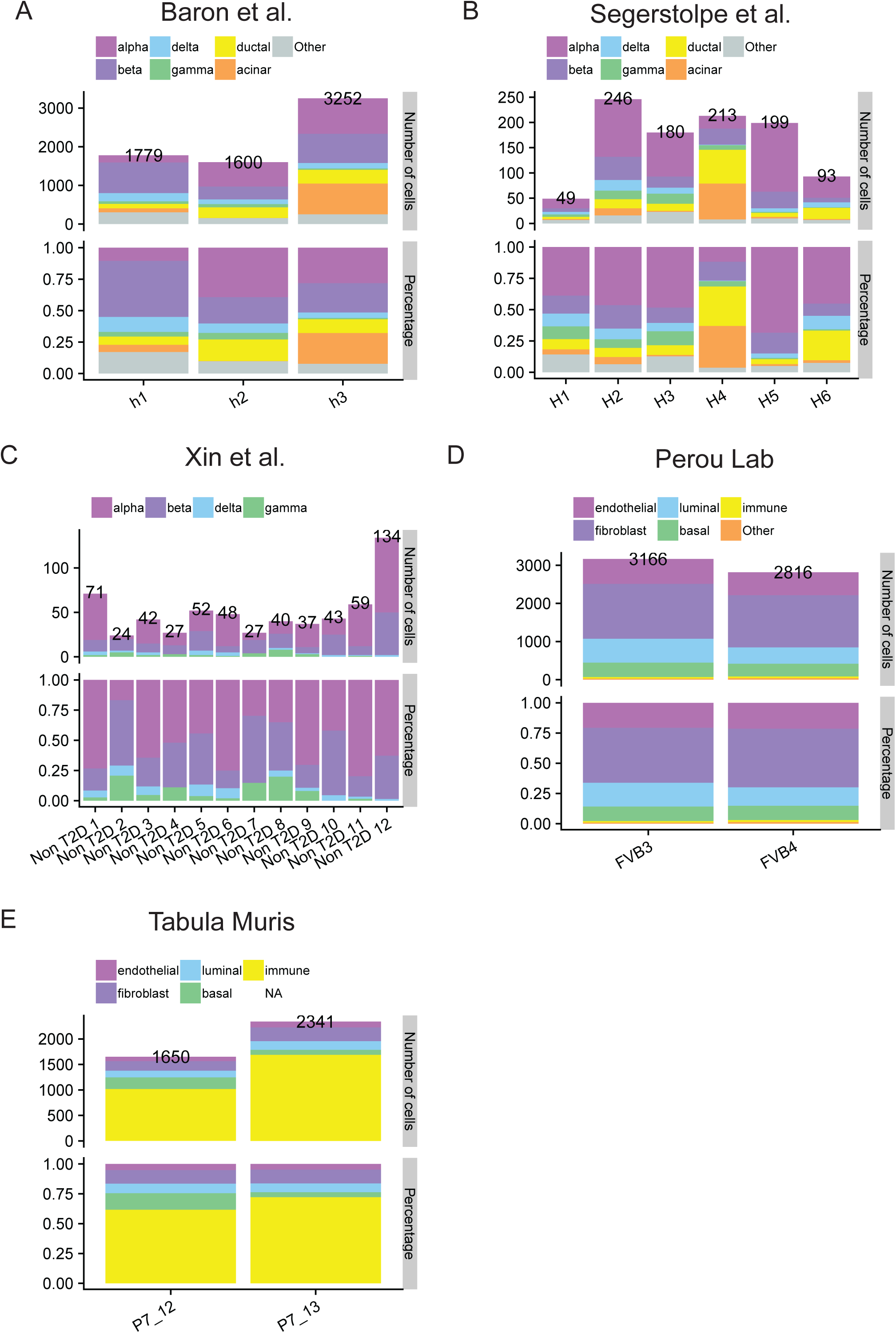
Number and percentage of single cells grouped by cell type clusters using scRNA-seq data of human pancreatic islets and mouse mammary glands. **A:** Baron et al. (2016). **B:** Asa Segerstolpe et al. (2016). **C:** Xin et al. (2016). **D:** Perou Lab. **E:** Tabula Muris.

**Table S1.** Benchmark of deconvolution results using simulated pseudo bulk samples of human pancreatic islets. The pseudo bulk samples were constructed by summing up raw read counts across all single cells from (A) Baron et al., (B) Segerstolpe et al., and (C) Xin et al.. The performance of deconvolution was assessed using measurements on the deconvolved and true cell-type proportions. SCDC outperforms Bseq-SC, and performs similar to MuSiC, when using only one reference set without ENSEMBLE, while naively pooled single cells without batch correction generally resulted in bad performance. With ENSEMBLE, SCDC performs similar to CIBERSORTx in two out of the three simulation setups, yet significantly better in the third, highlighting its performance stability. Clustering quality control (CQC) procedure resulted in improved deconvolution accuracy except for the cases involving Xin et al., which has a limited number of single cells per subject.

| Pseudo Bulk | Single-cell reference (number of cell types used) | Methods | mAD Before CQC | mAD After CQC | Pearson R Before CQC | Pearson R After CQC | $\Delta$ Pearson R |
| --- | --- | --- | --- | --- | --- | --- | --- |
| Baron et al. | Baron et al. (6 types) | MuSiC | 0.027 | 0.026 | 0.97 | 0.966 | 0.00 |
|  |  | Bseq-SC | 0.056 | 0.057 | 0.878 | 0.879 | 0.00 |
|  |  | Bisque | 0.021 | 0.021 | 0.982 | 0.98 | 0.00 |
|  |  | CIBERSORTx | 0.041 | 0.041 | 0.966 | 0.958 | -0.01 |
|  |  | **SCDC** | **0.029** | **0.03** | **0.961** | **0.96** | **0.00** |
|  | Segerstolpe et al. (6 types) | MuSiC | 0.056 | 0.049 | 0.892 | 0.9 | 0.01 |
|  |  | Bseq-SC | 0.073 | 0.08 | 0.808 | 0.789 | -0.02 |
|  |  | Bisque | 0.087 | 0.092 | 0.671 | 0.65 | -0.02 |
|  |  | CIBERSORTx | 0.084 | 0.076 | 0.852 | 0.87 | 0.02 |
|  |  | **SCDC** | **0.05** | **0.045** | **0.912** | **0.924** | **0.01** |
|  | Xin et al. (4 types) | MuSiC | 0.147 | 0.117 | 0.541 | 0.717 | 0.18 |
|  |  | Bseq-SC | 0.144 | 0.154 | 0.434 | 0.435 | 0.00 |
|  |  | Bisque* | 0.099 | 0.101 | 0.813 | 0.814 | 0.00 |
|  |  | CIBERSORTx | 0.190 | 0.198 | 0.324 | 0.230 | -0.09 |
|  |  | **SCDC** | **0.06** | **0.085** | **0.964** | **0.973** | **0.01** |
|  | Pooled single cells (6 types) | MuSiC | 0.127 | 0.123 | 0.391 | 0.418 | 0.03 |
|  |  | Bseq-SC | 0.108 | 0.108 | 0.694 | 0.698 | 0.00 |
|  |  | Bisque* | 0.06 | 0.061 | 0.849 | 0.841 | -0.01 |
|  |  | CIBERSORTx | 0.036 | 0.038 | 0.968 | 0.963 | -0.01 |
|  |  | **SCDC** | **0.117** | **0.108** | **0.493** | **0.607** | **0.11** |
|  | ENSEMBLE (6 types) | **SCDC** | **0.029** | **0.03** | **0.961** | **0.96** | **0.00** |

\*Bisque only used part of the subjects due to the encountered error when including subjects with a missing cell-type.

(B)
| Pseudo Bulk | Single-cell reference (number of cell types used) | Methods | mAD Before CQC | mAD After CQC | Pearson R Before CQC | Pearson R After CQC | $\Delta$ Pearson R |
| --- | --- | --- | --- | --- | --- | --- | --- |
| Segerstolpe et al. | Baron et al. (6 types) | MuSiC | 0.083 | 0.082 | 0.915 | 0.918 | 0.00 |
|  |  | Bseq-SC | 0.089 | 0.085 | 0.895 | 0.899 | 0.00 |
|  |  | Bisque | 0.094 | 0.096 | 0.642 | 0.627 | -0.02 |
|  |  | CIBERSORTx | 0.085 | 0.081 | 0.955 | 0.955 | 0.00 |
|  |  | **SCDC** | **0.078** | **0.078** | **0.922** | **0.924** | **0.00** |
|  | Segerstolpe et al. (6 types) | MuSiC | 0.029 | 0.027 | 0.968 | 0.975 | 0.01 |
|  |  | Bseq-SC | 0.064 | 0.06 | 0.875 | 0.898 | 0.02 |
|  |  | Bisque | 0.035 | 0.035 | 0.963 | 0.962 | 0.00 |
|  |  | CIBERSORTx | 0.06 | 0.061 | 0.949 | 0.951 | 0.00 |
|  |  | **SCDC** | **0.032** | **0.031** | **0.961** | **0.969** | **0.01** |
|  | Xin et al. (4 types) | MuSiC | 0.096 | 0.091 | 0.933 | 0.94 | 0.01 |
|  |  | Bseq-SC | 0.075 | 0.076 | 0.92 | 0.927 | 0.01 |
|  |  | Bisque* | 0.063 | 0.064 | 0.944 | 0.943 | 0.00 |
|  |  | CIBERSORTx | 0.083 | 0.084 | 0.910 | 0.901 | -0.01 |
|  |  | **SCDC** | **0.113** | **0.133** | **0.97** | **0.965** | **-0.01** |
|  | Pooled single cells (6 types) | MuSiC | 0.058 | 0.056 | 0.887 | 0.897 | 0.01 |
|  |  | Bseq-SC | 0.085 | 0.083 | 0.692 | 0.694 | 0.00 |
|  |  | Bisque* | 0.059 | 0.059 | 0.878 | 0.876 | 0.00 |
|  |  | CIBERSORTx | 0.071 | 0.069 | 0.967 | 0.968 | 0.00 |
|  |  | **SCDC** | **0.056** | **0.061** | **0.916** | **0.928** | **0.01** |
|  | ENSEMBLE (6 types) | **SCDC** | **0.032** | **0.031** | **0.961** | **0.969** | **0.01** |

\*Bisque only used part of the subjects due to the encountered error when including subjects with a missing cell-type.

(C)
| Pseudo Bulk | Single-cell reference (number of cell types used) | Methods | mAD Before CQC | mAD After CQC | Pearson R Before CQC | Pearson R After CQC | $\Delta$ Pearson R |
| --- | --- | --- | --- | --- | --- | --- | --- |
| Xin et al. | Baron et al. (4 types) | MuSiC | 0.189 | 0.188 | 0.71 | 0.711 | 0.00 |
|  |  | Bseq-SC | 0.185 | 0.182 | 0.72 | 0.723 | 0.00 |
|  |  | Bisque | 0.091 | 0.098 | 0.874 | 0.855 | -0.02 |
|  |  | CIBERSORTx | 0.180 | 0.177 | 0.733 | 0.736 | 0.00 |
|  |  | **SCDC** | **0.188** | **0.187** | **0.71** | **0.711** | **0.00** |
|  | Segerstolpe et al. (4 types) | MuSiC | 0.071 | 0.075 | 0.92 | 0.915 | -0.01 |
|  |  | Bseq-SC | 0.154 | 0.153 | 0.757 | 0.758 | 0.00 |
|  |  | Bisque | 0.078 | 0.076 | 0.902 | 0.897 | -0.01 |
|  |  | CIBERSORTx | 0.161 | 0.163 | 0.756 | 0.751 | -0.01 |
|  |  | **SCDC** | **0.067** | **0.072** | **0.929** | **0.925** | **0.00** |
|  | Xin et al. (4 types) | MuSiC | 0.035 | 0.035 | 0.976 | 0.976 | 0.00 |
|  |  | Bseq-SC | 0.066 | 0.062 | 0.894 | 0.906 | 0.01 |
|  |  | Bisque* | 0.066 | 0.066 | 0.924 | 0.925 | 0.00 |
|  |  | CIBERSORTx | 0.050 | 0.051 | 0.959 | 0.957 | 0.00 |
|  |  | **SCDC** | **0.072** | **0.081** | **0.938** | **0.925** | **-0.01** |
|  | Pooled single cells (4 types) | MuSiC | 0.061 | 0.068 | 0.919 | 0.906 | -0.01 |
|  |  | Bseq-SC | 0.117 | 0.117 | 0.827 | 0.828 | 0.00 |
|  |  | Bisque* | 0.062 | 0.063 | 0.942 | 0.938 | 0.00 |
|  |  | CIBERSORTx | 0.174 | 0.173 | 0.739 | 0.740 | 0.00 |
|  |  | **SCDC** | **0.064** | **0.086** | **0.916** | **0.901** | **-0.02** |
|  | ENSEMBLE (4 types) | **SCDC** | **0.072** | **0.081** | **0.938** | **0.925** | **-0.01** |

\*Bisque only used part of the subjects due to the encountered error when including subjects with a missing cell-type.

**Table S2.** Associating cell-type proportions with HbA1c levels in human pancreatic islet samples. A linear regression model (deconvolved cell-type proportion ∼ HbA1c + age + BMI + sex) is adopted for each cell type separately. SCDC through ENSEMBLE derived a p-value of 0.0019 for the association between the HbA1c levels and the beta cell proportions, more significant than those from deconvolution without ENSEMBLE.

| Cell type<br>(% As<br>Outcome) |  | Estimate | Std.<br>Error | P-value<br>using<br>ENSEMBLE | p-value<br>using<br>Baron et al. | p-value<br>using<br>Segerstolpe<br>et al. |
| --- | --- | --- | --- | --- | --- | --- |
| alpha | (Intercept) | 0.8079 | 0.2152 | 4.00E-04 | 1.00E-04 | 0.0398 |
|  | HbA1c | -0.0087 | 0.0287 | 0.7627 | 0.4346 | 0.6449 |
|  | age | -0.001 | 0.002 | 0.6248 | 0.7861 | 0.9251 |
|  | BMI | -0.0107 | 0.0082 | 0.1972 | 0.2033 | 0.7888 |
|  | sexFemale | 0.0457 | 0.0444 | 0.3069 | 0.4485 | 0.1597 |
| beta | (Intercept) | 0.4082 | 0.1051 | 2.00E-04 | 0.0316 | 0 |
|  | **HbA1c** | **-0.0452** | **0.014** | **0.0019** | **0.038** | **0.031** |
|  | age | 0.002 | 0.001 | 0.0484 | 0.2291 | 0.2332 |
|  | BMI | -0.0041 | 0.004 | 0.3115 | 0.6331 | 0.059 |
|  | sexFemale | -0.0628 | 0.0217 | 0.005 | 0.1952 | 0.004 |
| delta | (Intercept) | 0.053 | 0.0098 | 0 | 0 | 0 |
|  | HbA1c | -0.001 | 0.0013 | 0.4242 | 0.4243 | 0.4527 |
|  | age | -1.00E-04 | 1.00E-04 | 0.1188 | 0.1344 | 0.1256 |
|  | BMI | -0.0011 | 4.00E-04 | 0.0053 | 0.0058 | 0.0059 |
|  | sexFemale | 0 | 0.002 | 0.9983 | 0.8213 | 0.8473 |
| gamma | (Intercept) | 0.0112 | 0.0125 | 0.3719 | 0.3556 | 0.3827 |
|  | HbA1c | 0.0012 | 0.0017 | 0.483 | 0.5547 | 0.4622 |
|  | age | 1.00E-04 | 1.00E-04 | 0.336 | 0.2919 | 0.3508 |
|  | BMI | -7.00E-04 | 5.00E-04 | 0.137 | 0.1281 | 0.1467 |
|  | sexFemale | -0.0019 | 0.0026 | 0.4631 | 0.4248 | 0.4588 |
| acinar | (Intercept) | -0.0056 | 0.0777 | 0.9431 | 0.8045 | 0.4762 |
|  | HbA1c | 0.0202 | 0.0104 | 0.0549 | 0.0283 | 0.0517 |
|  | age | -0.0014 | 7.00E-04 | 0.0491 | 0.166 | 0.0476 |
|  | BMI | 0.0014 | 0.003 | 0.6332 | 0.7244 | 0.2173 |
|  | sexFemale | 0.0294 | 0.016 | 0.0706 | 0.0516 | 0.0574 |
| ductal | (Intercept) | -0.2747 | 0.1767 | 0.1244 | 0.249 | 0.1091 |
|  | HbA1c | 0.0335 | 0.0236 | 0.1597 | 0.2929 | 0.1213 |
|  | age | 5.00E-04 | 0.0016 | 0.7626 | 0.8249 | 0.5345 |
|  | BMI | 0.0152 | 0.0068 | 0.0278 | 0.0567 | 0.0513 |
|  | sexFemale | -0.0104 | 0.0365 | 0.7761 | 0.4099 | 0.9686 |

